# Xenogeneic Skin Transplantation Promotes Angiogenesis and Tissue Regeneration Through Vitamin D-Activated Trem2+ Macrophages

**DOI:** 10.1101/2021.02.26.432991

**Authors:** Dominic Henn, Kellen Chen, Tobias Fehlmann, Dharshan Sivaraj, Zeshaan N. Maan, Clark A. Bonham, Janos A. Barrera, Chyna J. Mays, Autumn H. Greco, Sylvia E. Moortgat Illouz, John Qian Lin, Deshka S. Foster, Jagannath Padmanabhan, Arash Momeni, Dung Nguyen, Derrick C. Wan, Ulrich Kneser, Michael Januszyk, Andreas Keller, Michael T. Longaker, Geoffrey C. Gurtner

## Abstract

Skin allo- and xenotransplantation are the standard treatment for major burns when donor sites for autografts are not available and have been shown to significantly accelerate wound healing. Although the cellular elements of foreign grafts are rejected, the extracellular matrix components integrate into the wound and may underlie their beneficial effects on wound healing. The molecular mechanisms defining the relationship between the immune response to foreign grafts and their impact on wound healing have not been fully elucidated. Here, we investigated changes in collagen architecture after xenogeneic implantation of clinically available human biologic scaffolds. We show that collagen deposition in response to the implantation of human split-thickness skin grafts (hSTSG) containing live cells recapitulates normal skin architecture, whereas human acellular dermal matrix (ADM) grafts led to highly aligned collagen deposition, characteristic of fibrosis and scar. Using single-cell RNA-sequencing, we show that macrophage differentiation in response to hSTSG is driven by vitamin D (VD) signaling toward Trem2+ subpopulations with an enrichment of pro-angiogenic and anti-fibrotic transcriptomic programs. We subsequently induced this regenerative subpopulation *in vitro* by treating bone marrow-derived cells with vitamin D3 and found that hydrogel delivery of Trem2+ macrophages significantly accelerated wound closure in a human-like murine excisional wound model. Our study identifies the preclinical therapeutic potential of Trem2+ macrophages to mitigate fibrosis and promote wound healing and provides a novel effective strategy to develop advanced cell therapies for complex wounds.

**One Sentence Summary:** Vitamin D-activated Trem2+ macrophages promote angiogenesis and mitigate fibrosis, providing a novel effective strategy to develop advanced cell therapies for complex wounds.

## INTRODUCTION

For over fifty years, transplantation of unmatched human skin grafts has been the clinical standard-of-care for major burn wounds when autologous skin grafts are not available due to the extent of the burn injury. Although the cellular elements of unmatched grafts are rejected by the host (*1*), extracellular matrix (ECM) elements integrate into the wound bed and are potentially beneficial to wound healing in patients with burns (*2–4*) and chronic wounds (*5, 6*). In mice, it has been shown that the implantation of xenogeneic dermal scaffolds does not induce a strong inflammatory response but has a strong impact on macrophage polarization, potentially inducing regenerative phenotypes (*7, 8*).

Macrophages are a heterogenous and highly plastic cell population and believed to be critical mediators of the cellular response during all stages of soft tissue injury. Traditional classification systems of macrophages describe “classically activated” M1 macrophages, dominating the acute inflammation stage after injury, and “alternatively activated” M2 macrophages, which are thought to be involved in reparative processes of soft tissue remodelling. These classifications, however, were derived from *in vitro* studies and have been shown to insufficiently characterize the broad spectrum of macrophage phenotypes found in physiologic and pathologic conditions *in vivo* (*9, 10*). Implantation of both synthetic and biological materials induces specific tissue microenvironments that likely drive macrophage subpopulations. Synthetic materials commonly instigate a pro-inflammatory response, whereas biological grafts can induce regenerative macrophage subpopulations (*7*), potentially suppressing the development of fibrosis by producing arginase 1 (ARG1), resistin-like molecule-α (RELMα), and interleukin-10 (IL-10) (*9, 11, 12*). To add to the complexity, other studies have suggested a pro-fibrotic role for “M2” macrophages due to their ability to stimulate fibroblast differentiation into myofibroblasts (*9, 13, 14*). Collectively, these findings indicate that the impact of macrophage heterogeneity on the process of wound repair and tissue fibrosis remains incompletely understood.

Here, we examine the impact of clinically used human biologic scaffolds on the innate immune response and wound healing over time in a murine model. Using single-cell RNA sequencing (scRNA-seq), we show that macrophage polarization in response to cellular human split-thickness skin grafts (hSTSG) is driven by vitamin-D (VD) signaling toward a distinct subpopulation characterized by a high expression of the lipid receptor *Trem2* (triggering receptor expressed on myeloid cells) as well as pro-angiogenic and anti-fibrotic gene expression profiles. We then confirmed that VD signaling drives clonal proliferation of myeloid cells during skin healing using myeloid cell-specific Cre-mediated recombination in a multi-color reporter mouse model. Finally, we developed an approach to induce regenerative Trem2+ transcriptomic programs *in vitro* by treating bone-marrow (BM) derived macrophages with vitamin-D3 (VD3,1,25-dihydroxycholecalciferol) and showed that Trem2+ macrophages significantly accelerate the healing of murine full-thickness excisional wounds.

## RESULTS

### Cellular human skin grafts remodel into a physiologic dermal collagen network after long-term implantation and attract regenerative macrophages

To characterize the functional role of macrophages in the cellular response to biologic scaffolds, we subcutaneously implanted either human split-thickness skin grafts (hSTSG) containing viable cells (*15*) or human-derived acellular dermal matrix (ADM) grafts into C57/BL6 (wild-type) mice (*8*). The grafts themselves and the overlying murine skin were explanted after 1, 3, 7, 14, and 28 days (n = 6-8 per group) (**Fig. 1A**). A sham surgery (creation of a subcutaneous pocket without graft implantation) served as a control to account for the impact of the surgical procedure itself (**fig. S1A**). We found that collagen deposition within both grafts significantly increased over time (P = 0.003), indicating that both scaffolds strongly influenced murine collagen production (**Fig. 1B** and **C**). To examine collagen architecture over time, we used the CT-FIRE algorithm to perform single fiber extraction and analysis in histologic images (*16*) as well as CurveAlign, a curvelet transform-based fibrillar collagen quantification platform to analyze fiber alignment as a surrogate parameter for fibrosis (*17*). While hSTSG and ADM both showed a significantly higher fiber alignment (*P < 0.05) compared to sham in the early implantation period (before day 7), the alignment of hSTSG decreased after day 7 and became similar to murine dermis on day 14, while ADM generated a consistently higher alignment than murine dermis throughout the entire implantation period until day 28. This chronic, increased alignment indicated progressive fibrosis following ADM implantation. In contrast, long-term implantation of hSTSG induced a basket-weave like fiber pattern resembling the physiologic dermal collagen architecture of murine skin, with a significantly lower alignment compared to ADM on day 28 (*P < 0.05) (**Fig. 1D,E, fig. S1B**). This was supported by principal component analysis (PCA) on all output parameters from CT-FIRE and CurveAlign, showing that hSTSG samples - but not ADM samples - clustered more closely together with samples from the sham surgery group (**Fig. 1F,** and **fig. S2).**

**Fig. 1.**
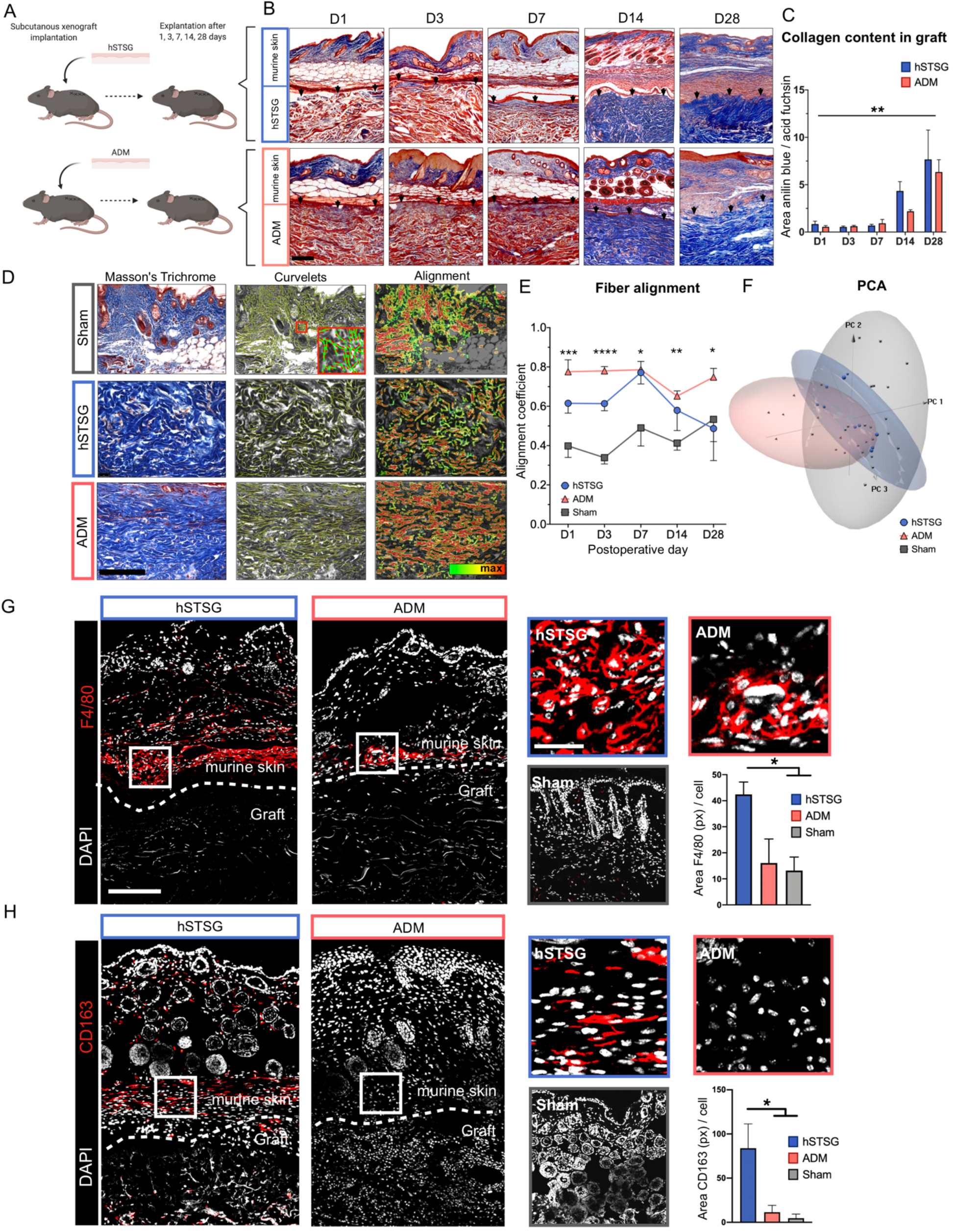
Human split thickness skin grafts remodel into a physiologic dermal collagen network after long-term *in vivo* implantation and attract regenerative macrophages. (**A**) Schematic of xenograft implantation model. Cryopreserved human split-thickness skin grafts (hSTSG) and human acellular dermal matrix (ADM) grafts were subcutaneously implanted into C57/BL6 mice. The grafts and the overlying murine skin were explanted after 1, 3, 7, 14, and 28 days (D1 – D28; n=6-8 per group). (**B, C**) Masson’s trichrome staining of tissue sections from the explanted grafts with overlying murine dermis showed a significant increase in collagen deposition within the graft over time (One-way ANOVA; **P = 0.006 for D14 vs. D1; *P = 0.03 for D14 vs. D3; *P = 0.02 for D14 vs. D7; **P = 0.007 for D28 vs. D1 and D3; **P = 0.005 for D28 vs. D7). Black arrows indicate the interface between graft and murine dermis. (**D**) Left panel: Masson’s trichrome staining of tissue sections showing collagen architecture of murine skin 28 days after sham surgery (top) as well as hSTSG (middle) and ADM (bottom) 28 days post implantation. Middle panel: CurveAlign output showing the curvelet transformation as an overlay on histologic image. Red dots indicate the center of fiber segments; green lines indicate the fiber orientation at that point. Small red square indicates the location of the magnified inset. Right panel: Heatmap of collagen fiber alignment. Red indicates the regions with the most aligned fiber angles. (**E**) Fiber alignment of ADM, hSTSG and murine dermis after sham surgery over time. One-way ANOVA: D1: ***P = 0.0009 for ADM vs. sham, **P = 0.02 for hSTSG vs. sham; D3: **P = 0.004 for ADM vs. hSTSG; ***P = 0.0002 for hSTSG vs. sham; ****P < 0.0001 for ADM vs. sham; D7: *P = 0.03 for hSTSG vs. sham; *P = 0.02 for ADM vs. sham; D14: **P = 0.001 for ADM vs. sham; D28: *P = 0.01 for ADM vs. sham. (**F**) Principal component analysis (PCA) on output parameters from CT-FIRE and CurveAlign (overall alignment, nearest neighbor alignment, nearest fiber distance, fiber width, fiber length) showed a high similarity between the fiber architecture of hSTSG (28 days after implantation) with sham surgery samples (all time points), whereas ADM samples clustered separately from sham surgery samples. (**G**) Immunofluorescent staining of explanted hSTSG and ADM with overlying murine skin (D14) as well as murine skin after sham surgery for the global macrophage marker F4/80 and (H) the regenerative macrophage marker CD163 (One-way ANOVA: F4/80: *P = 0.04, CD163: *P = 0.03 for hSTSG vs. ADM and sham). Dotted lines indicate the interface between murine dermis and implanted grafts. hSTSG = human split-thickness skin graft, ADM = acellular dermal matrix graft; white rectangles indicate the location of the magnified images of hSTSG and ADM (right panels); scale bars: 200µm in hSTSG and ADM overview images (left panels) and sham image, and 50µm in magnified hSTSG and ADM images (right panels).

We further performed fractal analysis to compare the histologic architecture between hSTSG and ADM grafts after implantation. Fractal analysis applies non-traditional mathematics to complex patterns that defy understanding with traditional Euclidean geometry (*18*). The overall tissue complexity of hSTSG, defined by the fractal dimension (FD), was significantly higher at D28 compared to ADM (*P = 0.04), which is in line with a more heterogenous basket-weave-like physiologic skin architecture (**fig. S3A** and **B**).

Since macrophages are the primary cell type that responds to the implantation of foreign bodies (*19*), we examined macrophage infiltration within the scaffolds and surrounding murine tissue on day 14, when the collagen architecture of the two grafts began to diverge. We observed a significantly higher number of macrophages around implanted hSTSG compared to ADM and sham by immunofluorescent staining for the generic macrophage marker F4/80 (*P=0.04, **Fig. 1G**) (*20*). Macrophages recruited to hSTSG also showed a significantly higher expression of the regenerative “M2” macrophage marker CD163 (*P = 0.03, **Fig. 1H**) (*21*), indicating that the hSTSG microenvironment might induce macrophage recruitment and polarization toward regenerative phenotypes.

### Human cellular elements degrade after murine subcutaneous implantation

To confirm that the identified cells were of murine and not human (i.e., grafted) origin, we performed real-time PCR (qPCR) for human Alu elements (primate-specific short interspersed elements [SINEs]), as previously described (*22*), on both non-implanted hSTSG and hSTSG explanted after 14 days of subcutaneous implantation. Murine skin was analyzed as a negative control and live human umbilical vein endothelial cells (HUVEC) were used as a positive control. High levels of human Alu DNA were detected in non-implanted hSTSG, which even exceeded those of HUVECs (*P < 0.01), indicating the presence of live human cells within hSTSG and confirming the findings from previous studies (*15*). After 14 days of subcutaneous implantation, Alu DNA levels in hSTSG had significantly decreased by several orders of magnitude, indicating that the human cells disappeared and only murine cells were present (**fig. S3C**).

### hSTSG promotes greater recruitment of circulating cells than ADM

To examine whether tissue resident or circulating cells were recruited to the grafts, we performed parabiosis of green fluorescent protein (GFP) expressing mice to wild-type (WT) mice and performed either hSTSG or ADM implantation or a sham surgery on the WT mice (**Fig. 2A**). Confocal laser scanning microscopy of the explanted grafts showed a significantly stronger recruitment of circulating GFP+ cells to hSTSG compared to ADM or sham tissue (*P = 0.01 vs ADM and **P=0.002 vs. sham, **Fig. 2B**). The circulating cells recruited to hSTSG showed a higher expression of vascular endothelial growth factor (VEGF) compared to those in ADM and sham tissue (VEGF area (px) per cell: *P = 0.04 for hSTSG vs. ADM and P = 0.06 vs. sham, **Fig. 2C**), suggesting that hSTSG implantation promotes pro-angiogenic cell populations. In accordance, we observed a significantly stronger vascularization of the murine tissue surrounding hSTSG compared to ADM and sham tissue (CD31 area (px) per cell: **P = 0.002 for hSTSG vs. ADM and ****P = 0.0006 vs. sham, **Fig. 2B**). Flow cytometry of cells isolated from explanted grafts confirmed a stronger infiltration of GFP+ cells in hSTSG vs. ADM (19.7% vs. 9.4% of all live cells) (**Fig. 2D, fig. S4**).

**Fig. 2.**
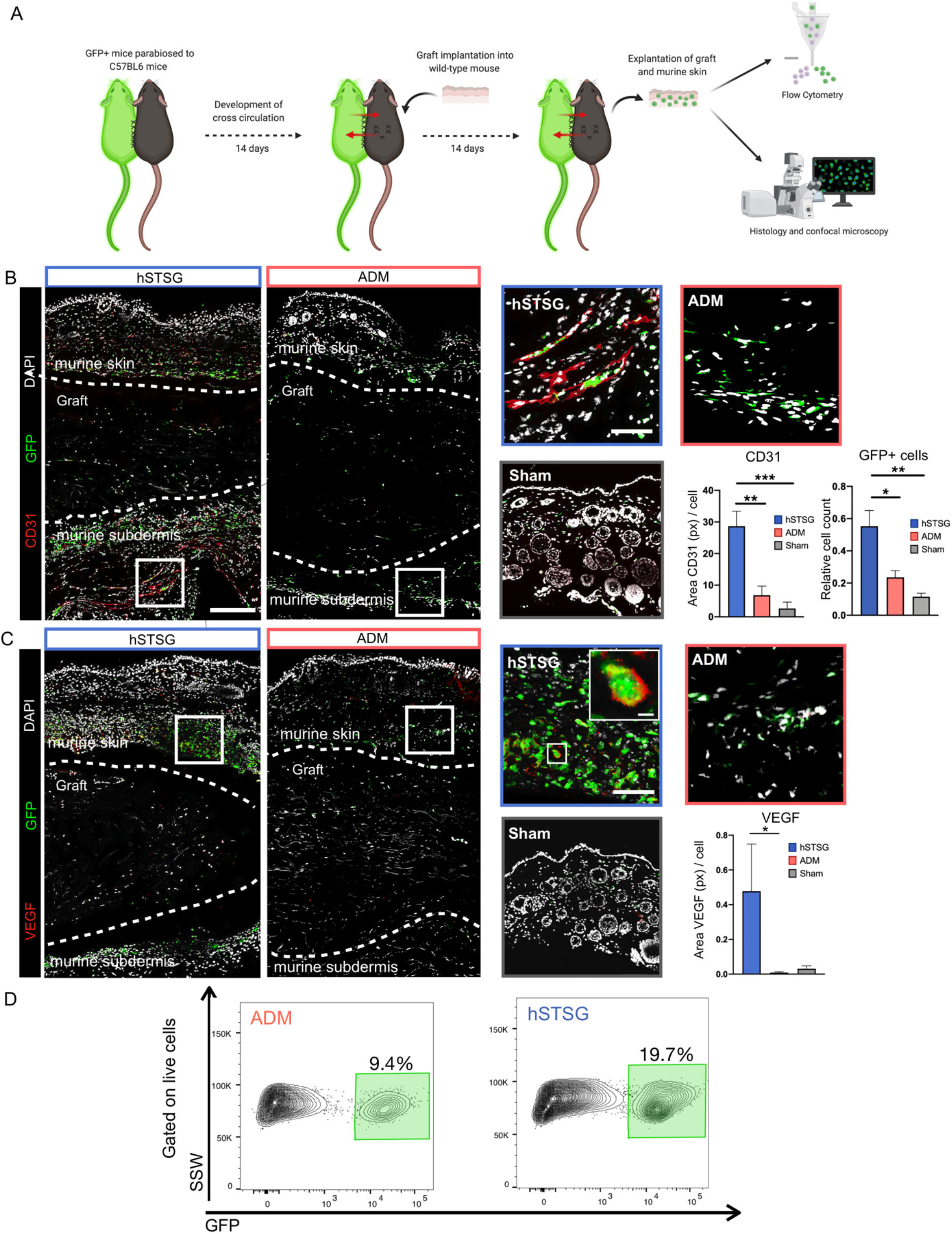
hSTSG promotes strong recruitment of circulating cells compared to ADM. (**A**) Schematic of the parabiosis model. Mice with generic expression of enhanced green fluorescent protein (GFP) were parabiosed to wild-type (WT) mice (C57BL/6). After 14 days and confirmation of cross-circulation, either hSTSG or ADM were subcutaneously implanted into the WT mice or a sham surgery was performed on the WT mice. The grafts were explanted after 14 days (n = 5 pairs per group). (**B**) Confocal laser scanning microscopy of cryo-sections of explanted grafts showed a higher proportion of circulating GFP+ cells in the murine tissue surrounding the graft compared to ADM and sham tissue (One-way ANOVA: relative cell count: *P = 0.01 vs. ADM, **P = 0.002 vs. sham). The murine tissue surrounding hSTSG showed a stronger vascularization compared to the tissue surrounding ADM and murine skin after sham surgery (One-way ANOVA: CD31 area in pixels (px) per cell: **P = 0.002 vs. ADM, ***P = 0.0006 vs. sham). (**C**) Circulating GFP+ cells recruited to hSTSG showed a higher expression of vascular endothelial growth factor (VEGF) compared to those in ADM and sham tissue (One-way ANOVA: VEGF area in px per cell: *P = 0.04 for hSTSG vs. ADM and P = 0.06 vs. sham). **(D)** Flow cytometry gating on live [4′,6-diamidino-2-phenylindole (DAPI)-negative] single cells showed a stronger GFP+ infiltration of hSTSG compared to ADM (19.7% vs. 9.4%) (n = 5 pairs per group). Dotted line on histology images indicates the interface between murine skin and graft. hSTSG = human split-thickness skin graft, ADM = acellular dermal matrix graft; white rectangles indicate the location of the magnified hSTSG and ADM images (right panels); scale bars: 200µm in hSTSG and ADM overview images (left panels) and sham image; 50µm in magnified hSTSG and ADM images (right panels); 20 µm in magnified image showing VEGF and GFP expressing circulating cell.

### Single-cell RNA sequencing and flow cytometry reveal heterogeneous macrophage subpopulations infiltrating hSTSG

To further understand the mechanisms driving these putatively regenerative cells, we performed scRNA-seq on hSTSG explanted on day 14 using the 10X Genomics Chromium platform (10X Genomics, Pleasanton, CA) (n = 5 mice, **Fig. 3A**). Data for 2,156 high-quality cells were embedded into UMAP-space using the Seurat package in R (*23, 24*) and identified as macrophages, monocytes, dendritic cells (DCs), granulocytes, T cells, fibroblasts, and endothelial cells (**Fig. 3B, fig. S5A**) (*25, 26*). We then further analyzed the largest cell population (1,108 cells), containing monocytes, macrophages, and DCs (**Fig. 3C** and **D**), as a subset and found 6 distinct cell clusters (labeled 0-5) (**Fig. 3E**). Among these, clusters 0, 2, and 3 were classified as macrophages, expressing the canonical markers *Adgre1* (F4/80) and *Cd68*; clusters 1 and 4 were identified as monocytes; and cluster 5 was classified as DCs (**Fig. 3F, fig. S5B-F**), We identified cluster defining marker genes and then confirmed them on the protein level using flow cytometric analysis of explanted grafts from a repeated group of mice (n = 5, **Fig. 3H, fig. S6A**). Cluster 0 was the largest macrophage cluster (31% of cells in the subset) and characterized by a strong expression of *Trem2* (triggering receptor expressed on myeloid cells 2), *Mertk* (proto-oncogene tyrosine-protein kinase MER), *Cd9, Cd63*, and a lack of *Cd163* expression (**Fig. 3F** and **G, fig. S5B** and **F**). Flow cytometry confirmed that F4/80+, MER+, CD163-cells were the most abundant macrophage subpopulation infiltrating hSTSG (37%) (**Fig 3H**). Cluster 1 cells were identified as Ly6c^high^ monocytes, showing an enrichment of *Ly6c2, Plac8*, *Cd86*, and MHCII encoding genes (*H2-Aa*, *H2-Ab1, H2-DMb1, H2-Eb1*), depletion of “M2” markers *Cd163, Mertk*, and *Cd209a* (**Fig. 3F** and **G, fig. S5C** and **F**), and confirmed on the protein level by FACS (26% F4/80+, MER-, CD163-, CD86+, CD209-cells) (**Fig. 3H**). Cluster 2 showed an enrichment of the regenerative macrophage markers *Mrc1* (CD206), *Cd163*, *Lyve1*, and *Retnla*, consistent with a recently described population of perivascular macrophages (**Fig. 3F** and **G, fig. S5D** and **F**) (*27*), and confirmed by FACS (F4/80+, Cd163+ cells) (**Fig. 3H**). Cluster 3 had the least distinct transcriptional profile and appeared to be a transitional state, showing an enrichment of the proteasome gene *Psmd13* and depletion of *Mertk* and *Cd86*; this profile was also confirmed with FACS (F4/80+, MER-, CD86-) (**Fig. 3F-H**). Cluster 4 appeared to be a precursor cell population with the potential to differentiate into monocytes or DCs and expressed the hematopoietic stem cell marker *Kit* (proto-oncogene tyrosine-protein kinase Kit), monocyte marker *Ly6c*, and DC lineage markers *Cd209a* (DC-SIGN) and *Flt3* (**Fig 3F** and **G, fig. S5E** and **F**). Cluster 5 was a transcriptionally distinct cluster of DCs, displaying a high expression of common DC markers *Ly75, Flt3*, and *Ccr7* (**Fig. 3F** and **G, fig. S5E** and **F**).

**Fig. 3.**
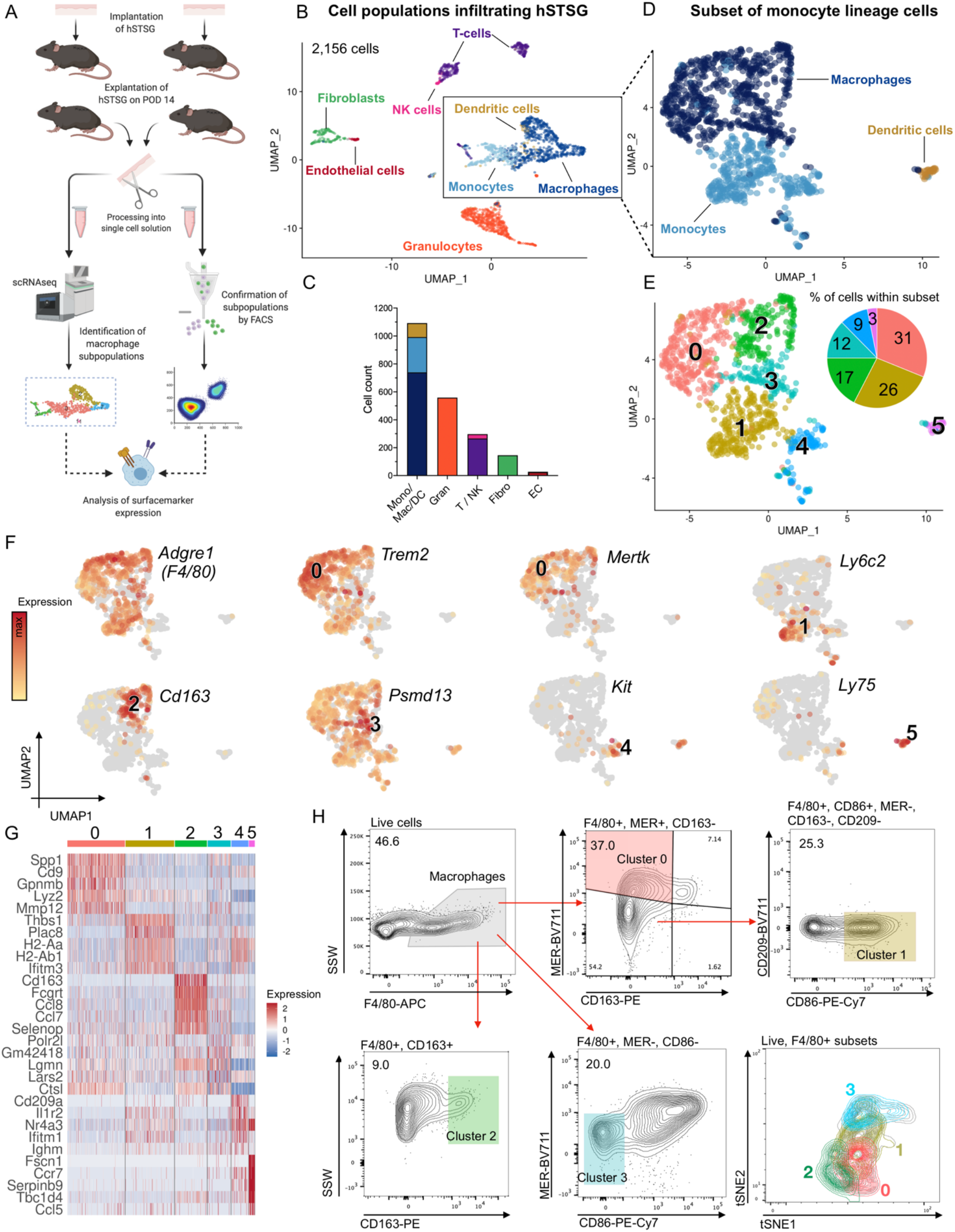
Single-cell RNA sequencing and flow cytometry reveal heterogeneous macrophage subpopulations infiltrating hSTSG. (**A**) Experimental overview. After 14 days of implantation, hSTSG were explanted (n = 5 mice), enzymatically digested, and processed for single-cell RNA-sequencing (scRNA-seq). Macrophage subpopulations identified by scRNA-seq were confirmed at the protein level in hSTSG explanted from a second group of mice (n = 5) using flow cytometry with a surface marker profile determined by the scRNA-seq analysis. (**B**) scRNA-seq yielded 2,156 high-quality cells. Cells (colored by cell type) were identified as macrophages, monocytes, dendritic cells (DCs), granulocytes, T cells, natural killer (NK) cells, fibroblasts, and endothelial cells. (**C**) Bar plot (colored by cell type) indicating the cell number within the populations of (B): mono = monocytes, macro = macrophages, gran = granulocytes, T / NK = T cells and NK cells, fibro = fibroblasts, EC = endothelial cells. (**D**) The largest population of 1,108 cells were identified as monocyte lineage cells (macrophages, monocytes, DCs). (**E**) The subset of monocyte lineage was composed of 6 distinct cell clusters (0 – 5). The pie chart (colored by cluster) shows the relative number of cells within the monocyte lineage subset. (**F**) Feature plots of both *Adgre1* (encodes for F4/80) as a global monocyte/macrophage marker as well as cluster defining markers. (**G**) Heatmap of the top 5 differentially expressed genes per cluster, sorted by average log fold-change and ordered by cluster number. (**H**) Flow cytometry gating strategy for cells isolated from hSTSG after 14 days of implantation (n = 5 mice). Single cells were gated for live (DAPI-negative) cells. Macrophages were identified by their expression of F4/80 based on the strong expression of *Adgre1* in monocytes and macrophages. Cluster defining expression of genes coding for surface markers was used to define a gating strategy for macrophage subpopulations. Cluster 1 was defined as F4/80+, MER+, CD163-. cluster 0: F4/80+, CD86+, MER-, CD163-, CD209-; cluster 2: F4/80+, CD163+; cluster 3: F4/80+, MER-, CD86-. Lower right panel: tSNE projection for virtual aggregate of flow cytometry data. Macrophage-subpopulations were back-gated and are colored by cluster color. hSTSG = human split-thickness skin graft.

### Trem2+ macrophages express pro-angiogenic and anti-fibrotic transcriptomic signatures

To further characterize the role of these myeloid subpopulations, we constructed a network of enriched gene sets from the Gene Ontology (GO) database (*28*). This analysis indicated that the macrophages in clusters 0 and 2 were enriched for gene sets associated with angiogenesis, cell migration, and epithelial cell proliferation, while the monocytes in cluster 1 were enriched for gene sets related to cytokine biosynthesis and metabolism (**Fig. 4A**). Gene sets associated with lipid metabolism were exclusively enriched in cluster 0, indicating that these cells may represent a recently defined subpopulation of lipid associated macrophages, which have shown anti-inflammatory properties and protective functions against adipocyte hypertrophy and metabolic disease (*29*) (**Fig. 4A**). Over-representation analysis (ORA) at the single cell-level using GeneTrail3 (*30*) confirmed a significant enrichment of gene sets for positive regulation of angiogenesis, wound healing, and collagen catabolism in the Trem2+ cells in cluster 0 (**Fig. 4B**), suggesting a pro-angiogenic and anti-fibrotic potential, corresponding with an overexpression of *Vegfa* and matrix-metalloproteinases (MMPs) such as *Mmp12* and *Mmp13* (**fig. S6B-D**). Summarizing these findings, we hypothesized that implantation of hSTSG stimulates macrophage polarization toward Trem2+ subpopulations that promote collagen remodeling and angiogenesis.

**Fig. 4.**
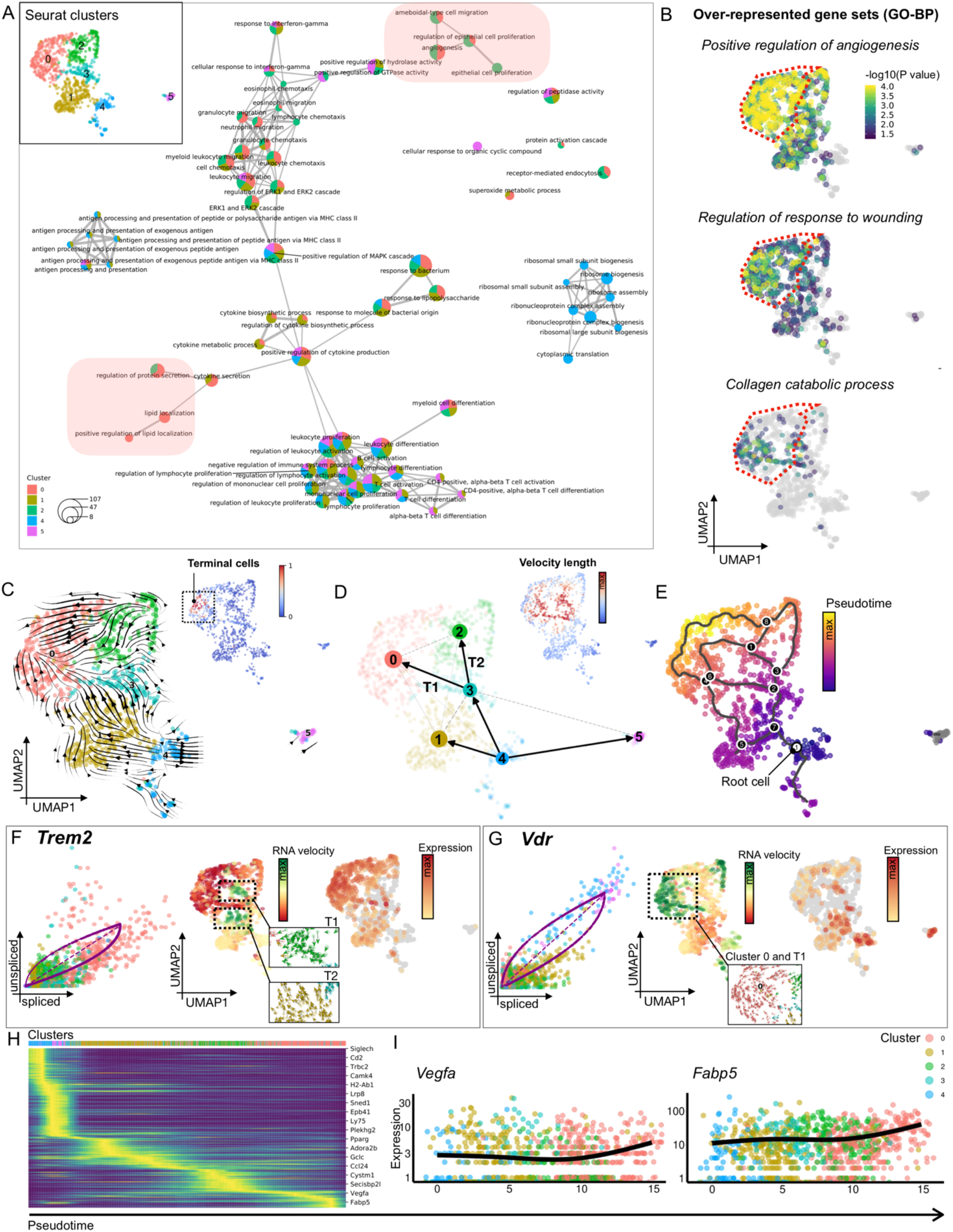
Trem2+ macrophages express pro-angiogenic and anti-fibrotic transcriptomic signatures. (**A**) Network diagram of enriched gene sets from the Gene Ontology (GO) database (biological process, BP). Each gene set is represented by a pie chart highlighting the relative contribution of each Seurat cluster. The size of the pie chart diagram represents the number of enriched genes per gene set. Gene sets enriched in clusters 0 and 2 that indicate the regenerative transcriptomic programs of these cells are highlighted in red. The Inset shows the Seurat clusters of the subset of Fig. 1E. (**B)** Over-representation analysis (ORA) of the indicated gene sets (GO-BP) projected on the UMAP embedding of monocyte lineage cells. Dotted lines indicate Seurat cluster 0, which showed the highest Trem2 expression. (**C**) RNA velocities derived from the dynamical model of scVelo and projected onto the UMAP embedding of the monocyte lineage subset. The main gene-averaged flow is visualized by velocity streamlines. Inset: terminal cell state located within Seurat cluster 0 as identified by scVelo. (**D)** Partition-based graph abstraction (PAGA) was used to quantify the connectivity of subpopulations which are projected onto the UMAP embedding with edge weights representing confidence in the presence of connections. T1 and T2 are the trajectories leading from cluster 3 to clusters 0 and 2. Inset: Cells colored by length of the velocity vectors as a correlate of the rate of differentiation. (**E)** Cells colored by pseudotime determined using Monocle3. Black nodes represent the branch points of the trajectories, white node represents the root node. (**F**) Left panel: Gene-resolved velocities for Trem2. The dotted line represents the estimated ‘steady-state’ ratio of unspliced to spliced mRNA abundance. RNA velocities are the residuals from the steady-state line, with positive velocities indicating an up-regulation of a gene, i.e., a higher abundance of unspliced mRNA than expected in the steady state. Middle panel: A high induction (positive RNA velocity) of Trem2 mRNA is found along the main differentiation trajectories T1 and T2 (highlighted by dotted lines with magnified images showing the velocity vectors of the trajectory cells). Right panel: Mature (spliced) mRNA expression of *Trem2* projected onto the UMAP embedding. (**G**) Left panel: Gene-resolved velocities for *Vdr* (coding for vitamin D receptor). Middle panel: A high induction (positive RNA velocity) of *Vdr* mRNA is found in cluster 0 (highlighted by dotted lines with magnified images showing the velocity vectors of cluster 0 cells). Right panel: Mature (spliced) mRNA expression of *Vdr* projected onto the UMAP embedding. (**H)** Heatmap highlighting genes with high correlation with velocity pseudotime. **(I)** Expression of *Vegfa* and *Fabp5* along pseudotime. Cells colored by Seurat cluster.

### Vitamin D receptor signaling drives transcriptional dynamics of macrophage differentiation

To investigate the transcriptional dynamics of macrophage differentiation, we performed RNA velocity analysis using scVelo (*31*). While comparison of mRNA abundance between cell clusters only captures a static snapshot, RNA velocity analysis predicts the future states of cellular subpopulations by integrating the relative abundance of nascent (unspliced) and mature (spliced) mRNA. Our analysis revealed a differentiation stream originating from cluster 4, which was identified as a root cluster, with a dual commitment toward DCs (cluster 5) and monocytes/macrophages (cluster 1 and 3), matching observations from our differential expression analysis (**Fig. 4C** and **D, fig. S6E-G**). The highest differentiation rate, indicated by the velocity vector length (**Fig 4D**, inset) was found along two major differentiation trajectories, T1 and T2, indicating that macrophage differentiation from the root cluster was driven along these paths toward the putative regenerative clusters 0 and 2 (**Fig. 4D**). The Trem2+ cells in cluster 0 were identified as the terminal differentiation state (**Fig 4C**, inset) and also demonstrated the most advanced progression along pseudotime from the root cells in cluster 4 (**Fig 4E, fig. S6H**) (*32*).

While cluster 0 showed the highest expression of mature, spliced Trem2 mRNA, a particularly strong expression of nascent, unspliced mRNA, characterized by positive velocities, was found along T1 and T2 (**Fig. 4F**), indicating an induction of Trem2 mRNA transcription. Interestingly, a strongly increased transcription rate for the vitamin D receptor encoding gene, *Vdr*, was found along T1 as well as in cluster 0, shown by an abundance of unspliced *Vdr* mRNA (**Fig. 4G**). This indicated that activation of the vitamin D receptor potentially drives T1, pointing to a critical importance of vitamin D signaling for the differentiation of macrophages into the regenerative Trem2+ subpopulation. Other co-regulated genes (modules) with a high correlation with pseudotime along the main differentiation trajectories included *Lpl* (lipoprotein lipase), a central regulator of lipid metabolism; *Mfge8* (milk fat globule-EGF factor 8 protein), which diminishes tissue fibrosis and promotes angiogenesis in cutaneous wound healing (*33, 34*); *Fabp5* (fatty acid-binding protein 5); and *Vegfa*. These genes are consistent with the transcriptional signature of Trem2+ lipid associated macrophages and demonstrate their pro-angiogenic and anti-fibrotic phenotypes (**Fig. 5H** and **I**).

**Fig. 5.**
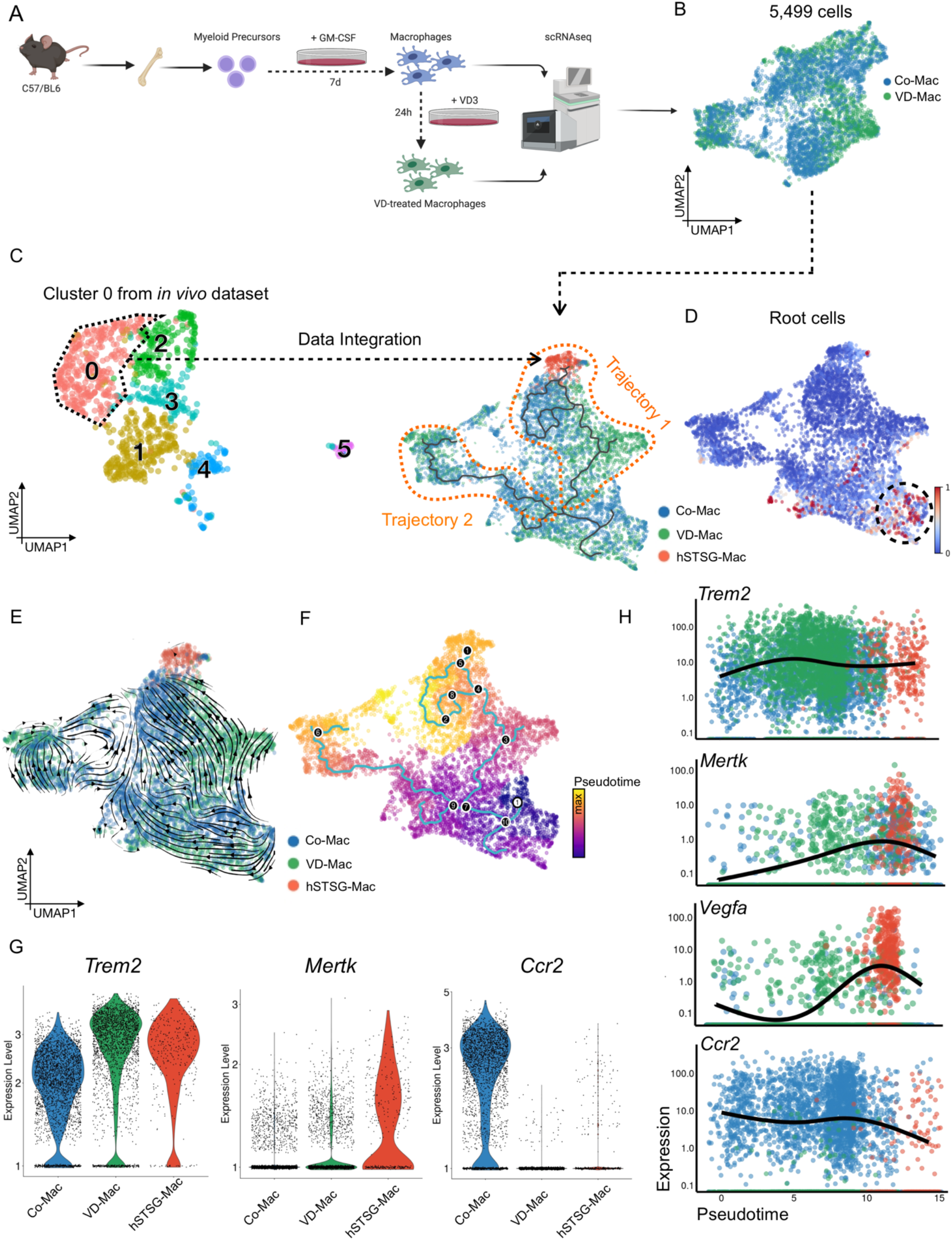
Vitamin D3 induces Trem2+ macrophage subpopulations *in vitro*. (**A**) Experimental workflow. Bone marrow (BM) was isolated from C57/BL6 mice and cells were differentiated into “M1” macrophages for 7 days in the presence of granulocyte-macrophage colony-stimulating factor (GM-CSF) according to standard protocols (Methods). The differentiated cells were then either stimulated with 1,25-dihydroxycholecalciferol (active form of vitamin D, VD3) for 24h or left untreated and both populations were analyzed using scRNA-seq. (**B**) UMAP embedding of cells colored by group. Co-Mac = control macrophages, VD-Mac = vitamin D3-treated macrophages. (**C**) Cells from cluster 0 from the *in vivo* monocyte-lineage subset (Fig 3E) were bioinformatically integrated with data from cultured macrophages. hSTSG-Mac = cluster 0 macrophages (Trem2+) from *in vivo* dataset. Trajectories (highlighted by orange dotted lines) were determined using Monocle3. (**D)** Root cells identified by RNA velocity analysis. **(E)** Main gene-averaged flow visualized by velocity streamlines and projected onto the UMAP embedding of the integrated dataset. **(F)** Cells colored by pseudotime using Monocle3. Black nodes represent the branch points of the trajectories, white node represents the root node. (**G**) Violin plots showing *Trem2, Mertk*, and *Ccr2* expression. Cells are colored by experimental group. (**H)** Expression of *Trem2*, *Mertk*, *Vegfa*, and *Ccr2* along pseudotime. Cells are colored by experimental group. hSTSG = human split-thickness skin graft.

### Vitamin D3 induces Trem2+ macrophage differentiation and clonal proliferation

To confirm that vitamin D receptor signaling drives macrophage differentiation toward Trem2+ subpopulations, we analyzed the impact of 1,25-dihydroxycholecalciferol (active form of vitamin D, VD3) on the transcriptional signatures of macrophages *in vitro*. Bone marrow cells were isolated from WT mice, differentiated into pro-inflammatory “M1” macrophages according to standard culture protocols (*35*), and either stimulated with VD3 for 24h or left untreated. The gene expression profiles of both populations were analyzed by scRNA-seq (**Fig. 5A** and **B**). VD3 treated cells showed a striking induction of *Vdr* (vitamin D receptor), *Cyp24a1* (encoding for VD3 24-hydroxylase, which catalyzes VD3 turnover), confirming a strong transcriptional response to VD3-treatment (**fig. S7A**). To relate these gene expression patterns to the Trem2+ macrophages from our *in vivo* dataset (cluster 0, **Fig. 3E**), we integrated both datasets (*36*) (**Fig. 5C, fig. S7B**) and observed that Trem2 expression in cultured macrophages treated with VD3 even exceeded *in vivo* levels, confirming our hypothesis that VD3 induces Trem2+ macrophage subpopulations (**Fig. 5G**). RNA velocity and pseudotime analysis identified an upward trajectory mainly containing VD3-treated cells (trajectory 1, **Fig. 5D-F**) and driven towards the *in vivo* Trem2+ macrophages (cluster i6 in integrated UMAP embedding, **fig. S7B**). This trajectory could represent the VD signaling-dependent trajectory leading to Trem2+ macrophage differentiation and was characterized by increasing *Mertk* and *Vegfa* with decreasing *Ccr2* expression, further confirming transcriptional programs observed *in vivo* (**Fig. 5G** and **H)**.

We next sought to investigate the effect of VD on myeloid cell proliferation during skin regeneration. We created stented full-thickness excisional wounds on the dorsum of Brainbow2.1-LyzMcre mice, in which *Cre* expression is driven by the myeloid cell specific *Lyz2* promotor and results in an irreversible recombination of the reporter locus, leading to stochastic expression of one of four fluorescent proteins (GFP, yellow fluorescent protein (YFP), red fluorescent protein (RFP), cyan fluorescent protein (CFP)) (**Fig 6A, fig. S8A, B**). Following daily VD3 injection into the wound bed, myeloid cells proliferated in a linear, polyclonal manner within the wound and generated a significantly higher number of clones compared to control wounds injected with a vehicle control (0.9% ethanol in PBS) and unwounded skin (***P = 0.0001 vs. control wound, ***P = 0.0002 vs. unwounded skin) (**Fig. 6B-E, fig. S9c**). Taken together, these findings show that VD signaling drives both myeloid differentiation into Trem2+ phenotypes as well as clonal myeloid cell proliferation during wound repair.

**Fig. 6.**
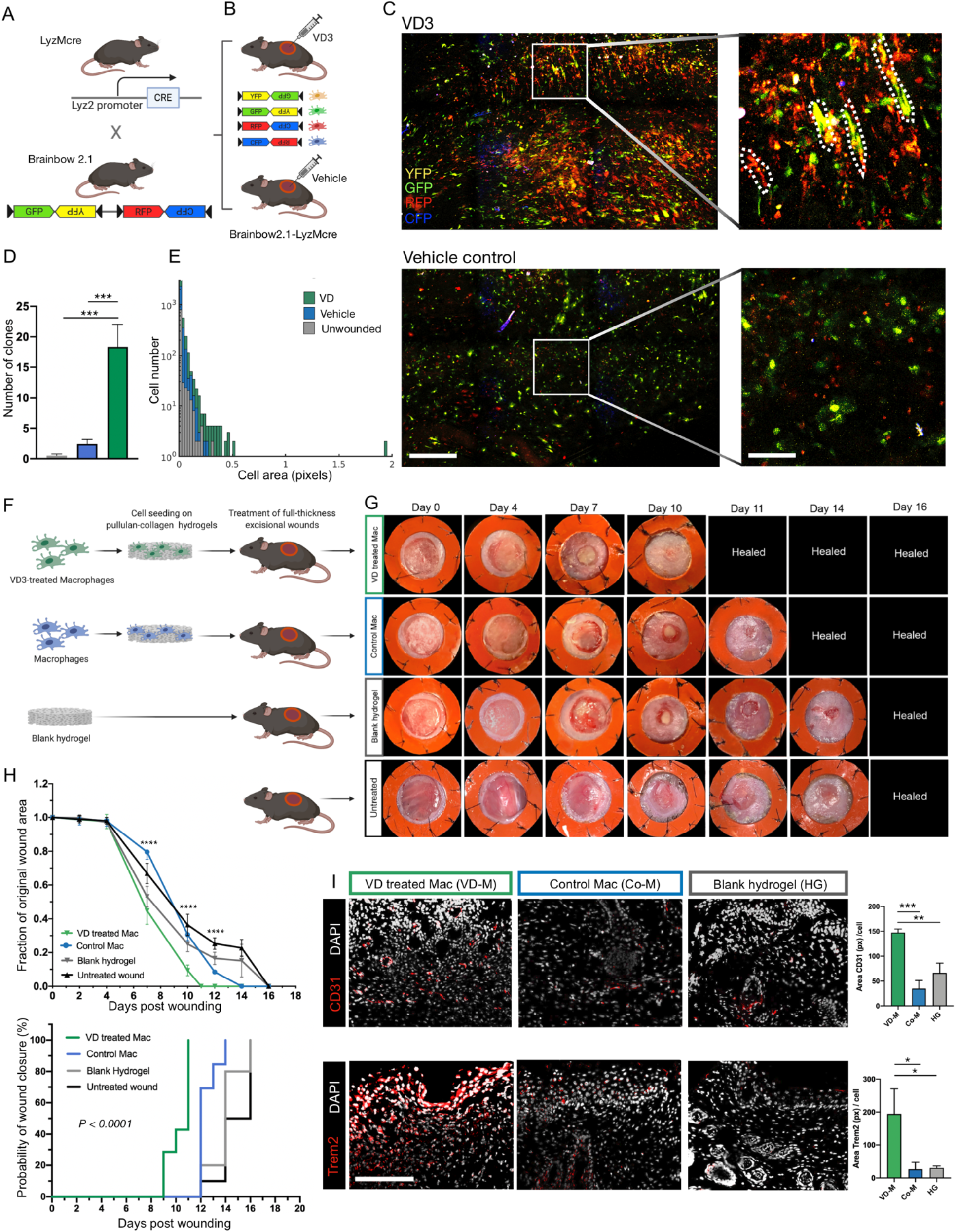
Trem2+ macrophage subpopulations accelerate healing of murine full-thickness excisional wounds. **(A)** Mice expressing Cre recombinase specifically in myeloid cells (Lyz2Mcre) were crossed with mice expressing a floxed transgenic four-color reporter construct (Brainbow 2.1) at the *Rosa26* locus (R26R-Brainbow 2.1). In the offspring, Cre expression is limited to myeloid cells expressing *Lyz2* and results in an irreversible recombination of the reporter locus, leading to stochastic expression of one of four fluorescent proteins (green fluorescent protein (GFP), yellow fluorescent protein (YFP), red fluorescent protein (RFP), cyan fluorescent protein (CFP)). **(B)** Stented full-thickness excisional wounds were created on the dorsum of *Brainbow2.1-LyzMcre* mice, and the wound bed was injected with either 1,25-dihydroxyvitamin D3 (VD3) or a vehicle control (0.9% ethanol in PBS) (n = 4 per group). **(C)** Myeloid cells were found to proliferate in a linear, polyclonal manner within the wound bed after VD3 injection (top panel), unlike wounds injected with the vehicle control (bottom panel) and unwounded skin (**fig. S9C)** which exhibited only minimal clonal proliferation; scale bars: 100 µm in overview, 20 µm in magnified images. **(D)** Cells formed a significantly higher number of clones after VD3 injection compared to wounds injected with the vehicle control or unwounded skin (One-way ANOVA: ***P = 0.0001 vs. control wound, ***P = 0.0002 vs. unwounded skin). **(E)** Myeloid cell clones within wound beds injected with VD3 showed a higher cell area compared to wounds injected with vehicle control or unwounded skin. **(F)** Two circular wounds of 6 mm diameter were created on the dorsum of C57/BL6 mice. The wounds were treated with pullulan-collagen hydrogels seeded with VD3-stimulated macrophages or untreated macrophages. Wounds in two control groups were either left untreated or were treated with unseeded hydrogels (n = 5 per group) (**G** and **H**). VD3-treated macrophages significantly accelerated wound healing and led to a significantly reduced wound area at day 7, 10, and 12 post-wounding compared to control macrophages, untreated hydrogels, and no treatment (Two-way ANOVA: day 7 and 10: ****P = 0.0001 vs. control macrophages, blank hydrogels and no treatment; day 12: ****P = 0.0001 vs. control macrophages and no treatment, *P = 0.02 vs. blank hydrogels). (**I**) Immunofluorescent staining of sections from healed wounds from mice treated with VD3-stimulated macrophages (VD-M, explanted on day 16 post-wounding) showed a significantly higher vascularization (CD 31area in pixels per cell: One-way ANOVA: ***P = 0.0002 for VD-M vs. Co-M, **P = 0.002 for VD-M vs HG) and Trem2 expression (Trem2 area in pixels per cell: One-way ANOVA: *P = 0.02 for VD-M vs. Co-M, **P = 0.03 for VD-M vs. HG) compared to wounds seeded with control macrophages (Co-M) or blank hydrogels (HG); scale bar: 200µm.

### Trem2+ macrophage subpopulations accelerate healing of murine full-thickness excisional wounds

To apply these findings toward clinically translatable outcomes, we investigated the capacity of VD3-induced Trem2+ macrophages to promote wound healing *in vivo*. We delivered BM-derived macrophages to stented full-thickness excisional wounds on WT mice using a soft pullulan-collagen hydrogel, previously developed by our group to effectively deliver cell therapies in both small and large animal wound models (*37–39*). Wounds were either left untreated, treated with blank hydrogels (HG) or treated with hydrogels seeded with VD3-stimulated macrophages (VD-M) or untreated control macrophages (Co-M) (**Fig. 6F**). VD-M significantly accelerated wound healing and led to a reduced wound area after day 7 compared to all other groups. Wounds treated with VD-M fully healed by day 11 (****P < 0.0001), whereas Co-M wounds healed by day 14 and HG treated wounds as well as untreated wounds healed by day 16 (**Fig. 6G** and **H**). Healed wounds treated with VD-M and explanted on day 16 showed significantly higher vascularization and Trem2 expression compared to tissue from both Co-M and HG treated wounds. These findings confirmed that Trem2+ macrophages induced by VD3 promote angiogenesis and accelerate wound healing (**Fig. 6I**).

## DISCUSSION

The innate immune system plays a critical role in the response to traumatic injury by clearing damaged cells and initiating the tissue repair process, which usually results in scar formation (*40–42*). Macrophages are a central component of the innate immune response and are critically involved in all phases of the injury response, from initial inflammation to tissue remodeling. Previous studies have investigated the innate immune response to foreign antigens as a first-line defense mechanism contributing to inflammation and graft rejection (*43, 44*). Accordingly, the beneficial effects of xenogeneic and allogeneic grafts on wound healing have so far been primarily attributed to their structural matrix component and not to cellular elements within the grafts (*2–4*). Our findings indicate that xenogeneic cells within transplanted grafts tune the innate immune response and induce reparative myeloid cells, leading to a beneficial remodeling of their collagenous matrix and surrounding host tissue that improves wound healing.

Using scRNA-seq, we showed that xenogeneic skin grafts induce myeloid cell differentiation toward reparative Trem2+ subpopulations. Trem2 is a transmembrane receptor of the immunoglobulin family, which binds anionic ligands such as phospholipids, sulfatides, and DNA. The disruption of cell membranes due to traumatic injury or apoptosis leads to an increase in free DNA and extracellular exposure of phospholipids, which could serve as a trigger for Trem2 receptor-ligand interactions (*45*). Using qPCR of primate specific DNA elements, we show that human cells within implanted xenogeneic grafts are cleared by the host over time. It seems likely that the resulting abundance of free foreign cellular elements provides a strong stimulus for macrophage differentiation into reparative Trem2+ phenotypes. The beneficial impact of local xenogeneic cell application via hSTSG on wound healing is in line with our previous work showing that systemic infusion of human MSCs promotes murine wound healing and macrophage polarization toward regenerative phenotypes within the wound (*46*). Our findings indicate that Trem2+ macrophages promote graft remodeling and breakdown of collagen fibers by secretion of MMPs, thus preventing fibrosis, and moreover, promote angiogenesis within the surrounding murine tissue. The observed regenerative effects resulting from degrading xenogeneic cells may be unique to skin transplantation and not translatable to xenotransplantation of other organs, which require long-term viability of donor cells to ensure functional organ integrity.

Multiple studies have reported on the anti-inflammatory properties of VD and its beneficial effect on wound healing (*47, 48*), and Vdr-KO models have highlighted the importance of VD on macrophage biology (*49*). However, the role of VD in macrophage differentiation has remained incompletely understood. We identified Trem2+ macrophages as previously unrecognized effector cells mediating the anti-inflammatory effect of VD3 during wound healing and fibrosis. Trem2+ macrophages have previously been shown to be involved in lipid metabolism and the regulation of hair follicle stem cell activity, pointing to an important role in skin regeneration (*50*). Variants in the Trem2 locus are associated with early-onset Alzheimer’s disease in humans, related to a reduced clearance of extracellular amyloid-β (Aβ)-containing plaques due to an impaired function of microglia within the brain (*51, 52*). In mouse models of Alzheimer’s disease, enhancement of Trem2 signaling using agonistic antibodies has recently shown beneficial effects on cognitive function through attenuation of neuroinflammation and improved plaque clearance within the brain (*53, 54*). The accelerated healing rates after enrichment of the wound bed with Trem2+ macrophages that we observed in our study might be due to a more efficient phagocytosis of dead cells and debris within the wound. Moreover, pro-angiogenic growth factors secreted by Trem2+ macrophages may improve vascularization and oxygen supply required for wound healing.

A limitation of our study is related to the fact that rodent models do not exactly recapitulate human wound healing; therefore future studies are needed to validate our findings in porcine wound models, which more closely mimic human biomechanics and skin physiology (*55, 56*). While our study demonstrates a strong regenerative effect of Trem2+ macrophages on wound healing, we have investigated only one dosing regimen of 500,000 cells applied weekly to the wound bed. Future studies investigating different dosages and dosing intervals are needed to maximize the therapeutic potential of our novel cell based therapy for wound healing before clinical translation can be achieved.

Our study uncovers the molecular background driving the long observed beneficial effects of unmatched skin transplantation on wound healing. To our knowledge, we are the first to demonstrate the preclinical therapeutic potential of Trem2+ macrophages to mitigate fibrosis and promote wound healing. We show that regenerative Trem2+ macrophages can be externally derived from bone marrow cell*s in vitro* using VD3 and then exogenously applied to wounds as a cell-based therapy.

Our approach to induce Trem2+ macrophages and deliver them to the wound bed using hydrogels provides a novel, effective strategy with a high potential for clinical translation. In the clinic, myeloid cells can be readily obtained in high quantities from the peripheral blood using clinically established cell apheresis and culture protocols (*57*), stimulated to differentiate into Trem2 phenotypes with VD3, and easily integrated with the current standard of care to improve wound healing after injury.

## MATERIALS AND METHODS

### Study design

The goal of our study was to investigate the impact of clinically used human biologic scaffolds on the innate immune response and soft tissue remodeling in a murine model over time. We chose a murine subcutaneous implantation model as this allowed us to interrogate the host response and specific microenvironment created by the implanted xenografts in a protected space without any potential external influence. scRNA-seq was employed to investigate the subpopulations of myeloid cells emerging after hSTSG implantation and the molecular mechanisms driving their differentiation. To confirm subpopulation defining markers on the protein level, we used flow cytometry on cells isolated from a repeated group of mice. The impact of VD signaling on clonal proliferation of myeloid cells during wound repair was confirmed using myeloid cell-specific Cre-mediated recombination in a multi-color reporter mouse model. To translate our findings for potential clinical applications, we developed an approach for induction of regenerative Trem2+ macrophages *in vitro* by treating bone-marrow (BM) derived macrophages with VD3 and seeded these cells on collagen-pullulan hydrogels. We have previously demonstrated that this technique preserves cell viability and enables efficient cell delivery to wounds (*38*). The capacity of Trem2+ macrophages to improve wound healing was investigated in a splinted full-thickness excisional wound model that limits the contraction of the murine panniculus carnosus muscle to recapitulate human physiological wound healing and allows murine wounds to heal by granulation tissue formation and re-epithelialization (*56*).

### Animals

All experiments were performed in accordance with Stanford University Institutional Animal Care and Use Committees and the NIH Guide for the Care and Use of Laboratory Animals. The study was approved by the Administrative Panel on Laboratory Animal Care (APLAC) at Stanford University (APLAC protocol number: 12080). The following mouse strains were obtained from the Jackson Laboratory (Bar Harbor, ME): C57BL/6J (wild-type), C57BL/6-Tg(CAG-EGFP)1Osb/J, Lyz2^tm1(cre)Ifo^/J (LyzMcre), and R26R-Brainbow 2.1. To breed Brainbow 2.1-LyzMcre mice, LyzMcre mice were crossed with R26R-Brainbow 2.1. mice. All animal surgeries were performed under inhalation anesthesia with isoflurane (Henry Schein Animal Health) at a concentration of 1-2% in oxygen at 3 L/min. The mice were placed in prone position, and dorsal fur was shaved. Skin was disinfected with betadine solution followed by 70% ethanol three times.

### Xenograft implantation model

For implantation of hSTSG and ADM, incisions of 1 cm were created on the dorsa of C57BL/6J (wild-type) mice (Jackson Laboratory, Bar Harbor, ME) and subcutaneous pockets were created by sharp dissection above the muscle fascia. hSTSG (TheraSkin®, Misonix, Inc, Farmingdale, NY) and ADM (AlloDerm, Allergan, Dublin, Ireland) were cut into 1 x 1 cm pieces and placed into the pockets, which were closed with 6-0 nylon sutures (Ethilon, Ethicon, Somerville, NJ). Sham surgeries were performed by creating pockets without graft implantation on a separate group of mice (n=6) as controls. The grafts and overlaying murine skin were explanted after 1, 3, 7,14, and 28 days and processed for analysis (n=6-8 per each group for ADM, hSTSG and sham surgeries)

### Parabiosis

Parabiosis was performed as previously described (*58*). Briefly, the corresponding flanks of GFP+ (C57BL/6-Tg(CAG-EGFP)1Osb/J) and C57BL6/J mice (WT) were shaved and disinfected with Betadine solution and 70% ethanol three times. Matching skin incisions were made from the olecranon to the knee joint of each mouse. The skin edges were undermined to create skin flaps of 1 cm width. 6-0 nylon sutures (Ethilon) were used to approximate the dorsal and ventral edges of the skin flaps and skin staples were used to close the longitudinal incisions. Buprenorphine was used for analgesia by subcutaneous injection every 8–12 h for 48 h after operation. Mice were monitored daily until the end of the experiment. After 14 days, peripheral blood chimerism was confirmed using fluorescent microscopy of the tail vein blood, before subcutaneous implantation of the grafts into the WT mice as described above. The grafts and the overlaying murine dermis were explanted from the parabiosed 14 days after implantation (n = 5 per group).

### Splinted Excisional Wound Model

Splinted full-thickness excisional wounds were created as previously described by *Galiano et al.* In this model, a silicone ring is sutured around the wound margin to stent the skin, mimicking human physiological wound healing by limiting the contraction of the murine panniculus carnosus muscle and allowing the wounds to heal by granulation tissue formation and re-epithelialization (*56*). Two full-thickness dermal wounds of 6 mm diameter were created on the dorsum of each mouse (C57BL/6J) using biopsy punches. A silicone ring was fixed to the dorsal skin using an adhesive glue (Vetbond, 3M, Saint Paul, MN) as well as 8 interrupted 6-0 nylon sutures placed around the outer edge of the ring to prevent wound contraction. The wounds were treated with collagen pullulan hydrogels or left untreated and covered with sterile dressings. Digital photographs were taken at the time of surgery and during every dressing change until the time of wound closure. For the hydrogel treatment study, wounds were harvested on day 16 post-wounding, when the wounds in all groups had healed.

Wounds in Brainbow 2.1-LyzM-Cre mice were injected daily with either 100 ng 1,25-dihydroxyvitamin D3 (dissolved in ethanol, and further diluted to 0.9% in PBS) or a vehicle control (0.9% ethanol in PBS). Wounds were harvested on day 5 for analysis of clonal macrophage proliferation (n = 5 per group).

### Histologic analysis of collagen content

Explanted tissue was fixed in 4% paraformaldehyde (PFA) in phosphate-buffered saline (PBS) for 24 hours. For Masson’s trichrome staining, tissue was dehydrated in a graded ethanol series and embedded into paraffin. Tissue sections were de-paraffinized and rehydrated through graded ethanol series. Masson’s trichrome staining (Sigma Aldrich) was performed according to the manufacturer’s recommendations and imaged using brightfield microscopy. We then implemented an algorithm in MATLAB to automatically deconvolve the color information of each Trichrome image (*59*). We determined a color matrix based on the stain specific RGB light absorption of these samples:

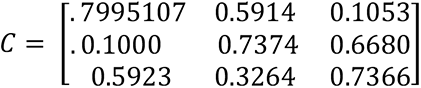

The top two rows correspond to stain specific RGB values of trichrome red and blue, respectively, and the columns represent the normalized vector values in the red, green, and blue channels. This algorithm allows for a robust and flexible method for objective immunohistochemical analysis of samples stained with up to three different colors.

### Immunofluorescent staining and confocal laser scanning microscopy

After fixation, tissue was dehydrated in 30% sucrose in PBS for at least 48 hours at 4°C. Tissue was then incubated in optimal cutting temperature compound (O.C.T., TissueTek, Sakura Finetek, Torrance, California) for 24 hours at 4°C and then cryoembedded in tissue molds on dry ice. Frozen sections were performed at 7 μm thickness on a cryostat. Antigen retrieval was performed using 0.01 M sodium citrate buffer in PBS (Abcam, Cambridge, MA), followed by blocking for 2 hours in 5% goat serum (Invitrogen, Waltham, Massachusetts) in PBS. Sections were then incubated in anti-F4/80 antibody (ab6640; Abcam), anti-CD31 antibody (ab28364; Abcam), anti-MMP13 antibody (ab39012; Abcam) or anti-VEGFA antibody (PA1-21796, ThermoFisher) over night at 4°C. Secondary antibodies were applied for 1 hour at room temperature (Goat anti-Rabbit IgG (H+L) Highly Cross-Adsorbed Secondary Antibody, Alexa Fluor Plus 488; Goat anti-Rat IgG (H+L) Cross-Adsorbed Secondary Antibody, Alexa Fluor 647; Goat anti-Rabbit IgG (H+L) Highly Cross-Adsorbed Secondary Antibody, Alexa Fluor Plus 647). Imaging was performed on a Zeiss LSM 880 confocal laser scanning microscope at the Cell Science and Imaging Facility at Stanford University. To obtain high resolution images of tissue samples, multiple images of 25x magnification were acquired using automatic tile scanning. Individual images were stitched together during acquisition using the ZEN Black software (Zeiss, Oberkochen, Germany).

### Quantification of immunofluorescent staining

Immunofluorescent staining was quantified using a code written in MATLAB adapted from previous image analysis studies by one of the authors (K.C.) (*60*). Briefly, confocal images were separated into their RGB channels and converted to binary to determine the area covered by each color channel. DAPI stain, corresponding to the blue channel, was converted to binary using *imbinarize* and an image-specific, automated threshold determined from the function *graythresh* in order to optimize the number of DAPI cells counted within the image. For the red and green channels, which corresponded to the actual protein stains, we converted these images to binary with a set a consistent threshold of ∼0.3 for all images of the same stain. This consistent threshold insured that our automated quantification of stain area would be unbiased. The area of red and green stains was then normalized by dividing by the number of cells, which we calculated as the number of DAPI nuclei above size threshold of 15 pixels. For clonal analysis, a size of 2000 pixels was chosen to delineate between either singular cells or clones. This threshold was chosen by visually inspecting images and counting an average size of inspected clones.

### Quantification of collagen fiber architecture

Analysis of fiber alignment of the murine dermis from the sham surgery group as well as the ADM and hSTSG grafts was performed on images (25X magnification) of Masson’s trichrome-stained tissue sections. Collagen fiber quantification was performed using CT-FIRE and CurveAlign, an open-source software package for automatic segmentation and quantification of individual collagen fiber (http://loci.wisc.edu/software/ctfire) (*16*). Briefly, CurveAlign quantifies all fiber angles and strength of alignment within an image, while CT-FIRE analyzes individual fiber metrics such as length, width, angle, and curvature. It can also extract other variables such as localized fiber density and the spatial relationship between fiber and the associated boundary. The average fiber parameters for each mouse were used for statistical analysis. Statistical analysis was performed in Prism8 (GraphPad, San Diego, California) using Student’s t test or one-way analysis of variance (ANOVA) with Tukey’s multiple comparisons test. The Rayleigh test was used to assess the circular distribution of the calculated vectors of alignment. Data are presented as means ± SEM. P values <.05 were considered statistically significant.

PCA was performed with centering to mean zero and scaling to unit variance using the prcomp function in R. Plots were generated with 95% confidence intervals using the ggbiplot and pca3d packages.

### Flow cytometry

Analysis of murine cells isolated from explanted grafts was performed according to published protocols for flow cytometry on murine tissue (*61*). In brief, hSTSG and ADM were explanted on day 14 from the parabiosis and non-parabiosis implantation models, then micro-dissected and incubated in serum-free Dulbecco’s Modified Eagle Medium (DMEM) with 240 U of collagenase IV/ml for 1h at 37°C in a rotating oven. Digested tissue was filtered, centrifuged and stained with BV711 rat anti-mouse Mer antibody, PE/Cy7 rat anti-mouse CD86 antibody, PE rat anti-mouse CD163 antibody, BV605 rat Anti-Mouse CD209a antibody, and APC rat anti-mouse F4/80 antibody (Biolegend, San Diego, CA). 4′,6-diamidino-2-phenylindole (DAPI) was used to stain dead cells. Flow cytometry was performed on a BD FACS Aria (Becton Dickinson, San Jose, CA) and data was analyzed using FlowJo (Becton Dickinson, San Jose, CA).

### Realtime PCR (qPCR)

Alu-qPCR was performed using a protocol and primers (101 F and 206 R) previously developed by Funakoshi et al. (*22*). The total reaction volume was 20 μl and contained 10 μl of TaqMan Universal Master Mix II, no UNG (Thermo Fisher Scientific), 0.2 μM forward and reverse primers, 0.25 μM hydrolysis probe, and the appropriate amount of genomic DNA (40 ng per reaction), on an Applied Biosystems 7900 instrument (Thermo Fisher Scientific). qPCR conditions were 1 cycle of 95 °C for 10 min, followed by 50 cycles of 95°C for 15s, 56°C for 30s and 72°C for 30s. All samples were run as triplicates. Beta actin was used as a reference gene. Fold-changes were calculated using the 2^-ΔΔ^method (*62*).

qPCR for genotype confirmation of transgenic Brainbow 2.1-Lyz2M-Cre mice was performed from DNA isolated from tail tissue by Transnetyx (Cordova, TN).

### Single cell RNA-seq data processing, normalization, and cell cluster identification

Base calls were converted to reads using the Cell Ranger (10X Genomics, Pleasanton, CA, USA; version 3.1) implementation *mkfastq* and then aligned against the GRCh38 v3.0.0 (human) genome using Cell Ranger’s count function with SC3Pv3 chemistry and 5,000 expected cells per sample. Cell barcodes representative of quality cells were delineated from barcodes of apoptotic cells or background RNA based on a threshold of having at least 300 unique transcripts profiled, less than 100,000 total transcripts, and less than 10% of their transcriptome of mitochondrial origin. Unique molecular identifiers (UMIs) from each cell barcode were retained for all downstream analysis. Raw UMI counts were normalized with a scale factor of 10,000 UMIs per cell and subsequently natural log transformed with a pseudocount of 1 using the R package Seurat (version 3.1.1)(*24*). Aggregated data were then evaluated using uniform manifold approximation and projection (UMAP) analysis over the first 15 principal components(*23*). Cell annotations were ascribed using the SingleR package (version 3.11) against the ImmGen database (*25, 26*) and confirmed by expression analysis of specific cell type markers. Louvain clustering was performed using a resolution of 0.5 and 15 nearest neighbors.

### Generation of characteristic subpopulation markers, enrichment and over-representation analysis

Cell-type markers were generated using Seurat’s native *FindMarkers* function with a log fold change threshold of 0.25 using the ROC to assign predictive power to each gene. The 50 most highly ranked genes from this analysis for each cluster were used to perform over-representation analysis (ORA) using clusterProfiler (version 3.1.1). Using GeneTrail 3 an ORA was performed for each cell using the 500 most expressed protein coding genes on the gene sets of the Gene Ontology(*30*). P-values were adjusted using the Benjamini-Hochberg procedure and gene sets were required to have between 2 and 1000 genes. Stacked violin plots were generated using the Scanpy package (*63*).

### Pseudotime analysis

Pseudotime analysis was performed using the Monocle 3 package in R (version 3 0.2.0)(*64*). A principal graph was learned from the reduced dimension space using reversed graph embedding with default parameters. The principalGraphTest function using the Moran’s *I* statistic was employed to identify correlated genes on trajectory embedded in the manifold. As indicated by RNA velocity analysis, cluster 4 was chosen as a root of the trajectory.

### RNA velocity analysis

RNA velocity analysis was performed using scVelo (*31*). In contrast to previous steady-state models, which falsely assume that all genes share a common splicing rate, scVelo uses a likelihood-based dynamical model to solve the full transcriptional dynamics of splicing kinetics. Thereby RNA velocity analysis can be adapted to transient cell states and heterogeneous cellular subpopulations as in our dataset. Partition-based graph abstraction (PAGA) was performed using the sc.tl.paga function in scVelo. To find genes with differentially regulated transcriptional dynamics compared to all other clusters a Welch t-test with overestimated variance to be conservative was applied, using the sc.tl.rank_velocity_genes function. Genes were ranked by their likelihood obtained from the dynamical model grouped by Seurat clusters. A pseudotime heatmap was created by plotting the expression of the genes identified by sc.tl.rank_velocity_genes along velocity-inferred pseudotime. The terminal transcriptional states were identified as end points of the velocity-inferred Markov diffusion process.

### Cultivation of bone-marrow derived macrophages and cell seeding

Bone marrow isolation and cultivation of cells into macrophages was performed according to previously published protocols (*65, 66*). Briefly, femurs and tibias were removed from C57/BL6 mice after euthanasia. The epiphyses on both ends of the long bones were removed. The bones were flushed with 5 ml RPMI to collect the bone marrow. The epiphyses were crushed using a pestle and mortar to isolate the bone marrow. Red blood cells were lysed with ACK lysis buffer. Cells were strained and washing in complete media 2×10^6^ per 100 mm dish in 10 ml RPMI containing 10% fetal bovine serum (FBS), 100 U/ml penicillin, 0.1 mg/ml streptomycin, 2 mM glutamine, 50 μM 2-mercaptoethanol, and 200 murine granulocyte-macrophage colony-stimulating factor (GM-CSF, Peprotech, Frankfurt, Germany). On day 3 10 ml complete media were added to the cultures and on day 6 10 ml of media were collected, cells were spun down and re-suspended in fresh media. On day 7, half of the cultures were stimulated with 10^-6^M 1,25 dihydroxy-cholecalciferol (Sigma-Aldrich) for 24 h. Cells were collected and seeded on pullulan-collagen hydrogels with a diameter of 8 mm at a concentration of 500,000 cells per hydrogel and wound, using a previously described capillary force seeding technique (*38*). We have previously demonstrated that this approach preserves cell viability as well as hydrogel architecture and enables efficient cell delivery (*38*).

### Statistica Analysis

Data are shown as mean ± SEM and were analyzed using Prism 8 (GraphPad, La Jolla, CA). Two-group comparisons were performed with Student’s t*-*test (unpaired and two-tailed). One- or two-way ANOVA (analysis of variance), followed by Benajmini Hochberg post hoc test were performed for comparisons of > 2 groups as appropriate. P *<* 0.05 was considered statistically significant. The statistical methods used for scRNA-seq analysis are described in the specific sections above.

## Supplementary Materials

**Fig. S1.**
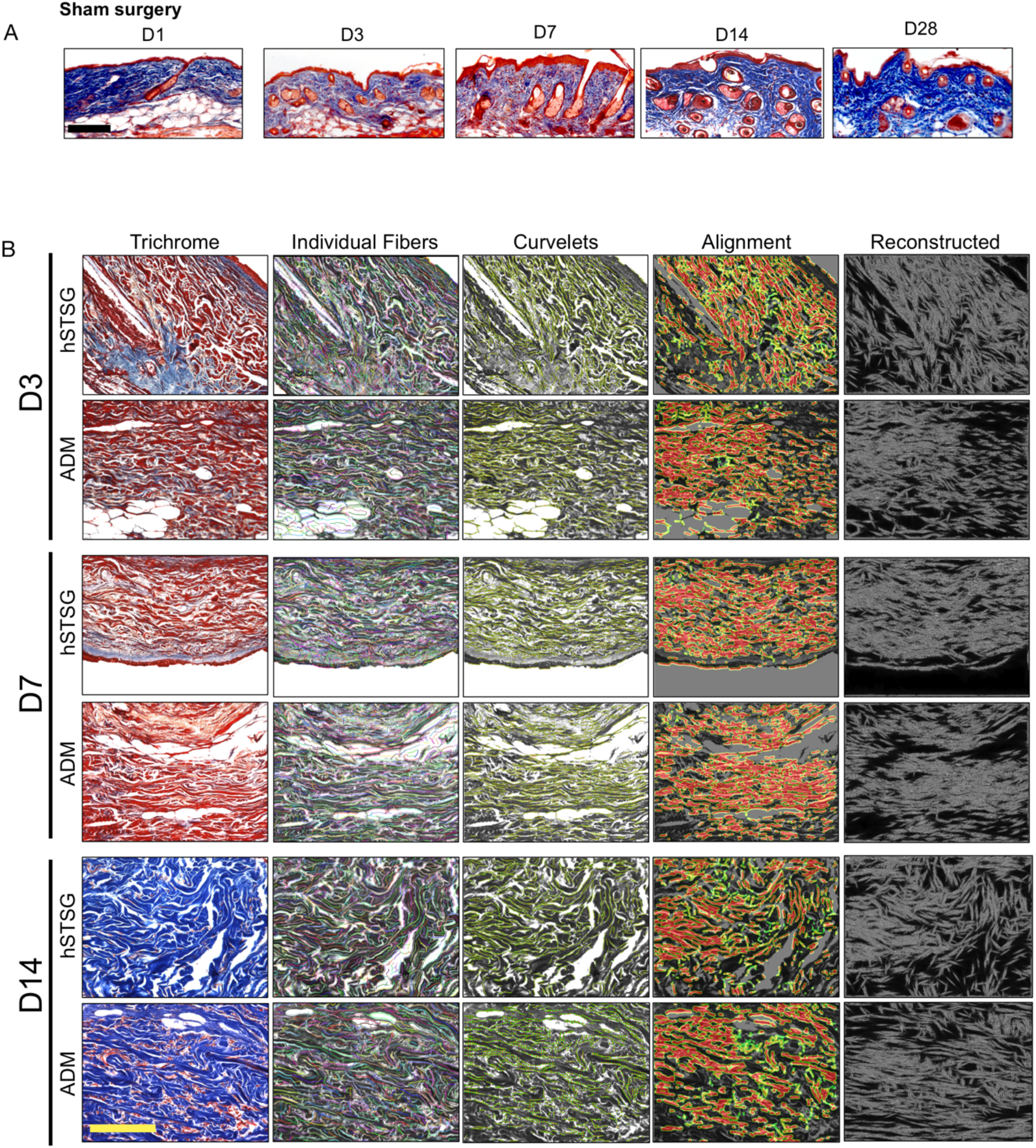
Sham group histology and collagen fiber analysis over time. (**A**) Masson’s trichrome staining of tissue sections from the sham group explanted on days 1, 3, 7, 14, and 28 post-surgery (D1 – D28). **(B)** Far left: Magnified image of collagen architecture of hSTSG and ADM (Masson’s trichrome staining). Left: individual fibers identified with CT-FIRE. Middle: curvelet transformation as overlay on histologic image, red dots to indicate the center of fiber segments, green lines indicate the fiber orientation at that point. Right: heatmap of alignment with red indicating the regions with the most aligned fiber angles; far right: image reconstructed using CurveAlign; scale bars: 200µm. hSTSG = human split-thickness skin graft, ADM = acellular dermal matrix graft.

**Fig. S2.**
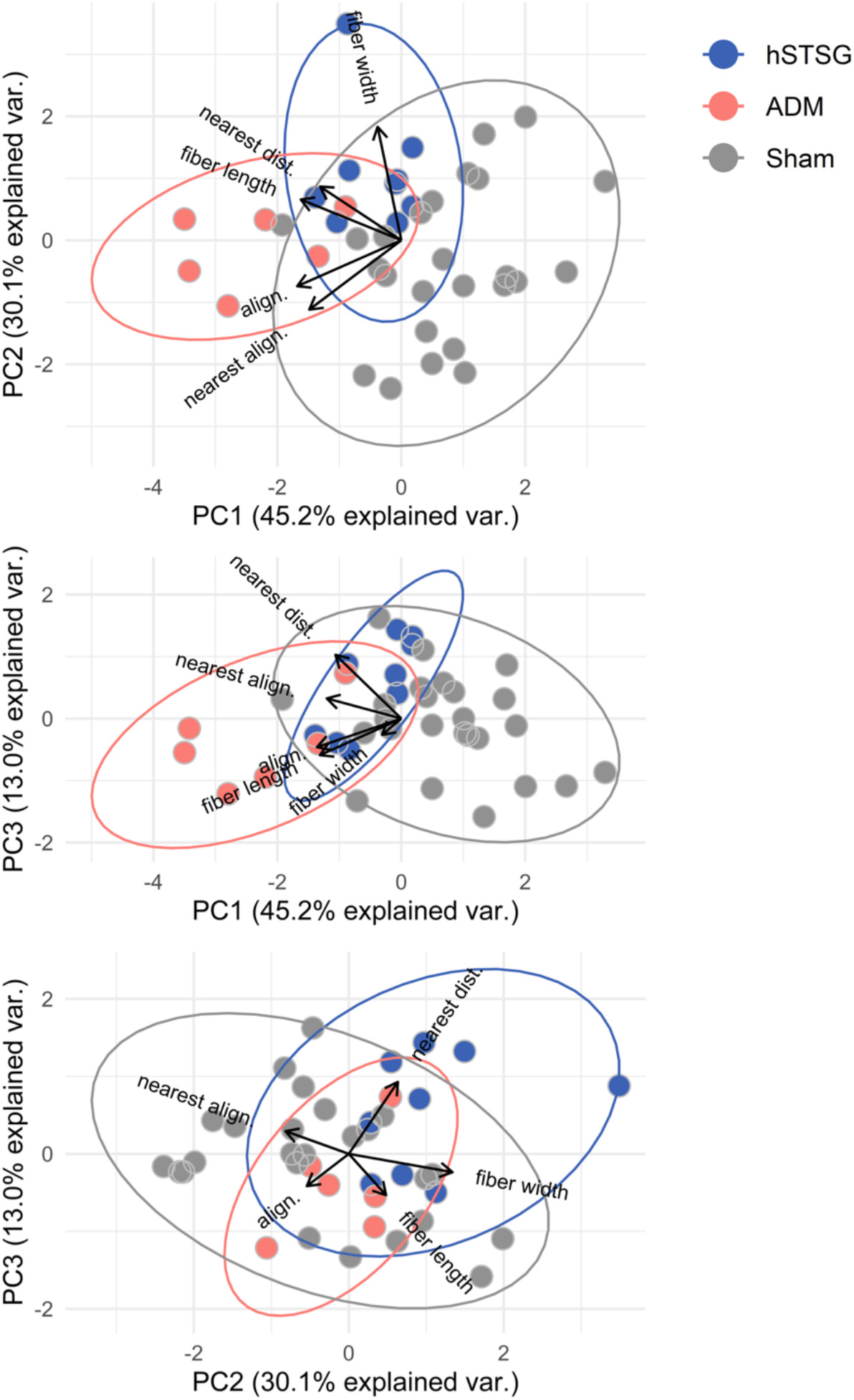
Principal component analysis (PCA) on CT-FIRE and CurveAlign output. Principal component analysis (PCA) on output parameters from CT-FIRE and CurveAlign analysis of collagen fibers from hSTSG, ADM (28 days post implantation) and skin mice with sham surgery 28 days post surgery. align = alignment, dist = distance, hSTSG = human split-thickness skin graft, ADM = acellular dermal matrix graft.

**Fig. S3.**
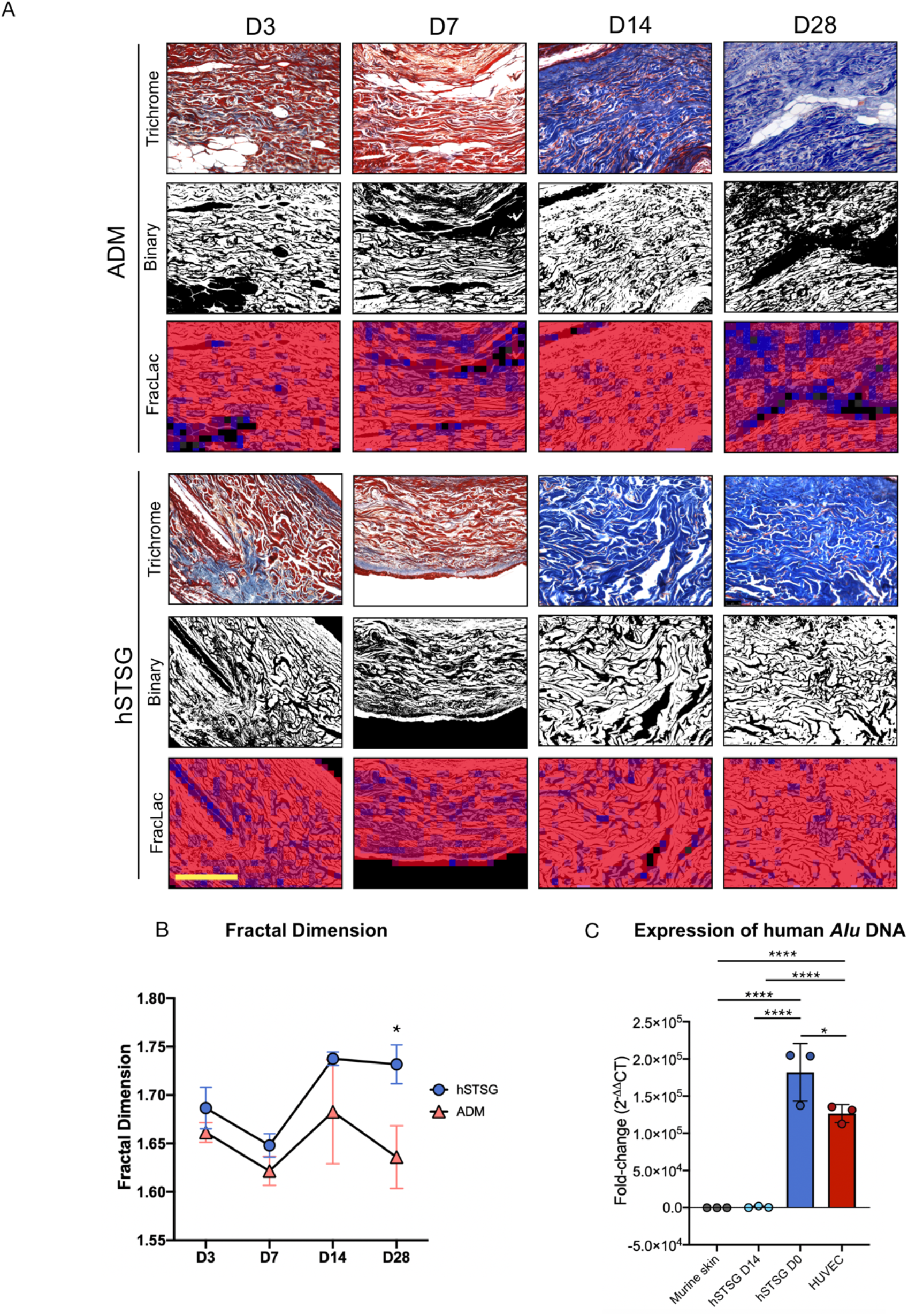
Fractal analysis and qPCR for human Alu elements. (**A** and **B**) Fractal analysis comparing the histologic architecture between hSTSG and ADM. Two-way ANOVA (n = 5 per group): *P= 0.04; scale bar: 200 µm. (**C**) Real-time PCR (qPCR) for human Alu elements in non-implanted hSTSG (hSTSG D0) and hSTSG explanted after 14 days of subcutaneous implantation into wild-type mice (hSTSG D14). Native murine skin was analyzed as a negative control and live human umbilical vein endothelial cells (HUVEC) were used as a positive control. The HUVEC group represents 3 different cell culture plates. All samples were analyzed as triplicates. Data is shown as fold-change computed by 2^-ΔΔ^method (*62*); One-way ANOVA (n = 3 per group): * P = 0.01; **** P < 0.0001, hSTSG = human split-thickness skin graft, ADM = acellular dermal matrix graft.

**Fig. S4.**
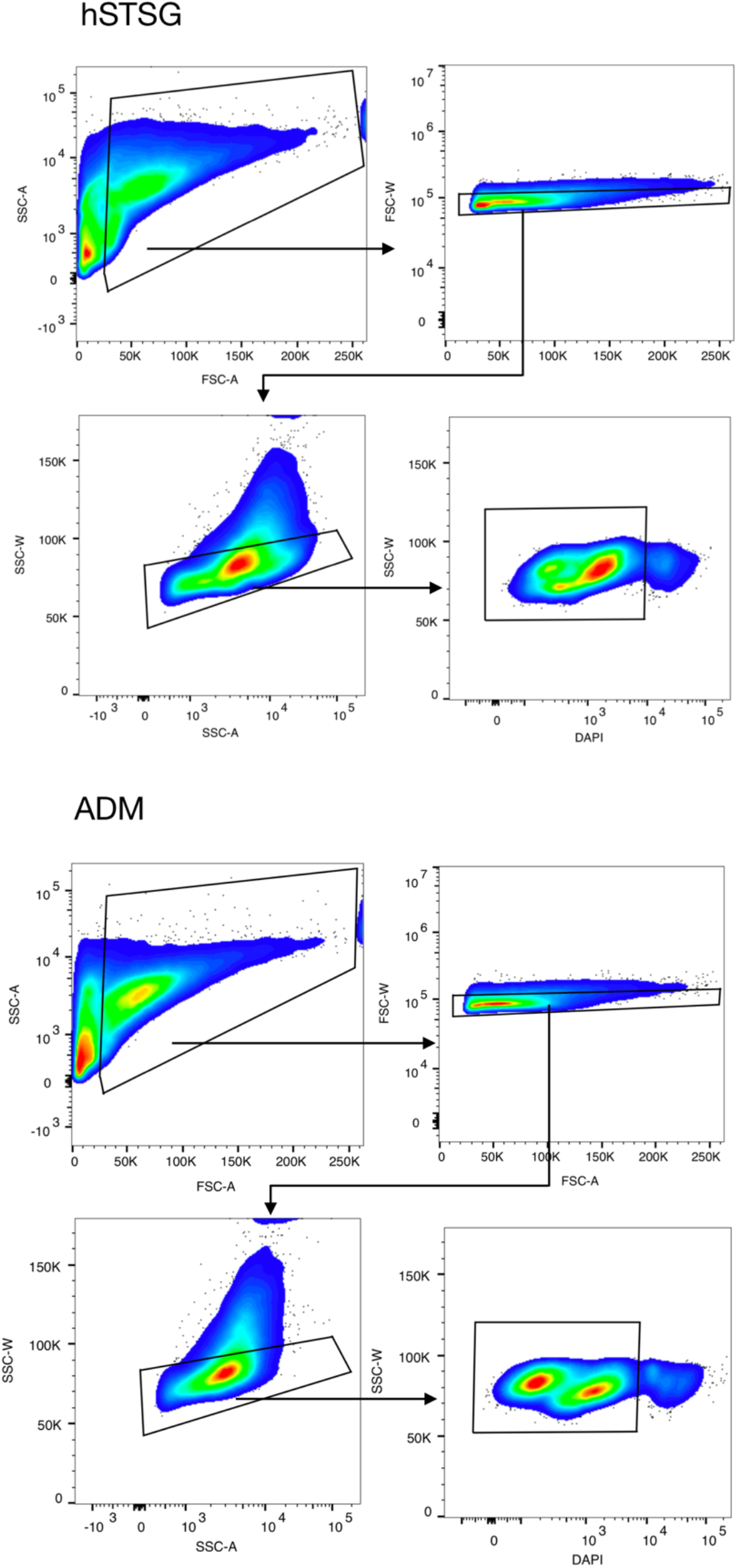
Flow cytometry pre-gating on cells from parabiosis model. Pre-gating strategy for flow cytometry on cells isolated from grafts explanted from parabiosis model (n = 5 pairs per group). hSTSG = human split-thickness skin graft, ADM = acellular dermal matrix graft.

**Fig. S5.**
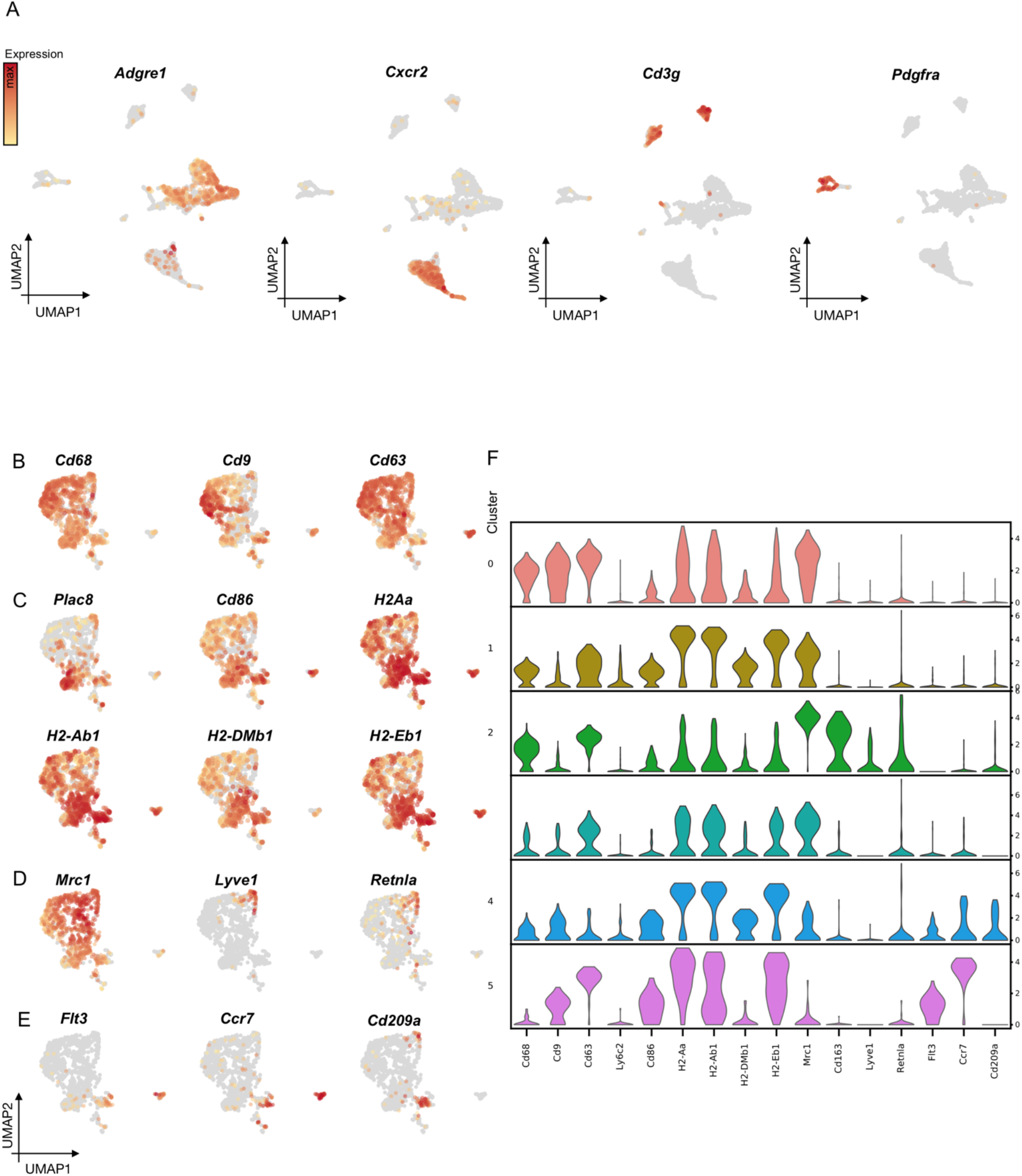
Cluster defining marker genes in full scRNA-seq dataset and myeloid cell subcluster. (**A**) Expression of the characteristic cell type marker genes, *Adgre1* (macrophages), *Cxc*r2 (neutrophilic granulocytes)*, Cd3g* (T cells), *Pdgfra* (fibroblasts) projected onto the UMAP embedding. Expression of the characteristic marker genes of myeloid cell subclusters: **(B**) cluster 0, (**C**) cluster 1, (**D**) cluster 2, (**E**) cluster 4 and 5 projected onto the UMAP embedding and (**F**) shown as stacked violin plot.

**Fig. S6.**
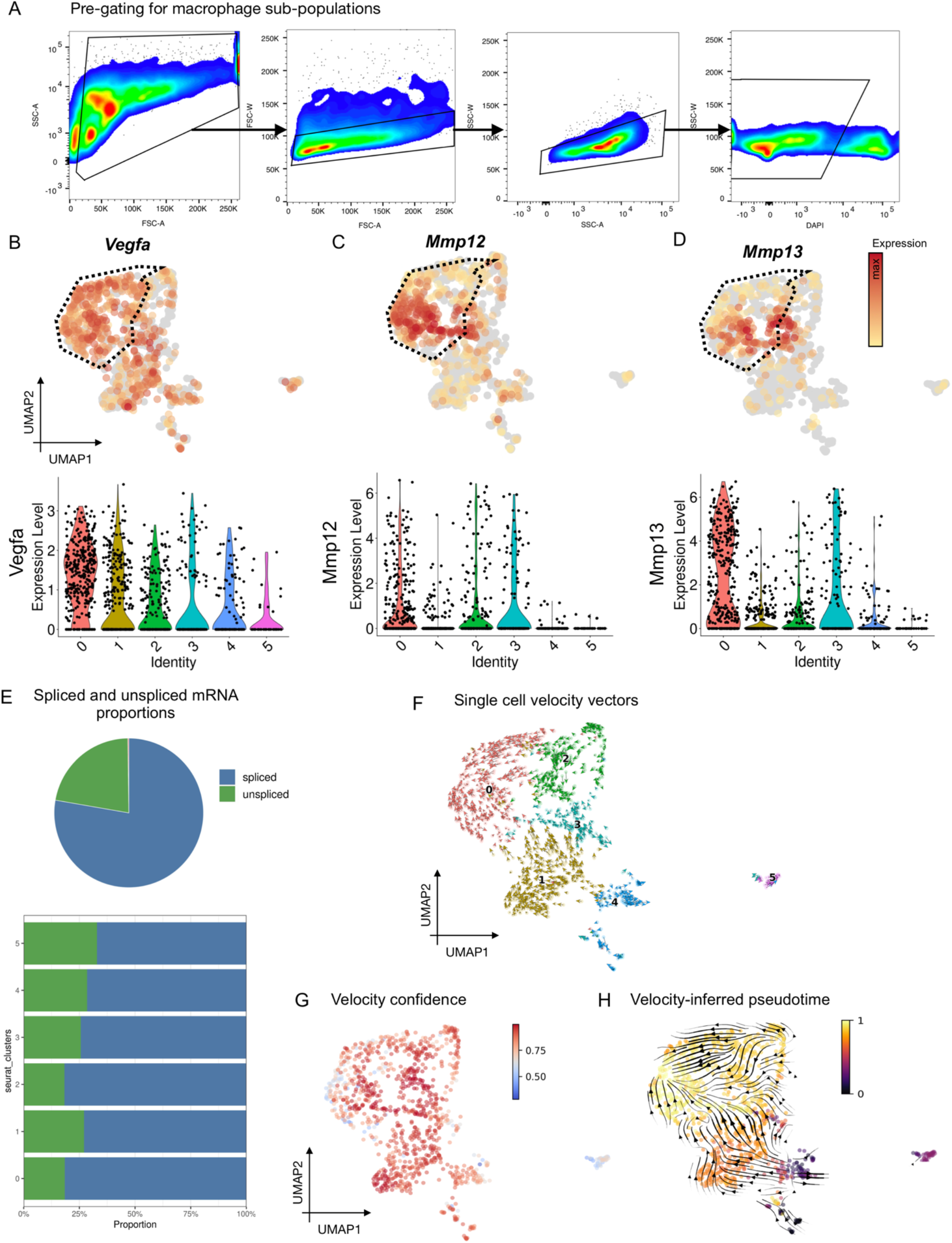
Flow cytometry pre-gating on macrophage subpopulations, scRNAseq analysis. (**A**) Flow cytometry pre-gating strategy for macrophage subpopulations to confirm clusters identified by single-cell RNAseq. (**B**) Expression of *Vegfa* (**C**), *Mmp12*, (**D**) *Mmp13* projected onto UMAP embedding of myeloid cell subset (top row) and shown as violin plot of the 6 clusters (bottom row). (**E**) Ratio of unspliced (green) and spliced (blue) mRNA within the myeloid cell subset (pie chart) and within each myeloid cell cluster, as determined by scVelo. (**F**) Single cell velocity vectors. (**G**) Confidence of RNA velocity analysis. (**H**) Velocity-inferred pseudotime determined by scVelo.

**Fig. S7.**
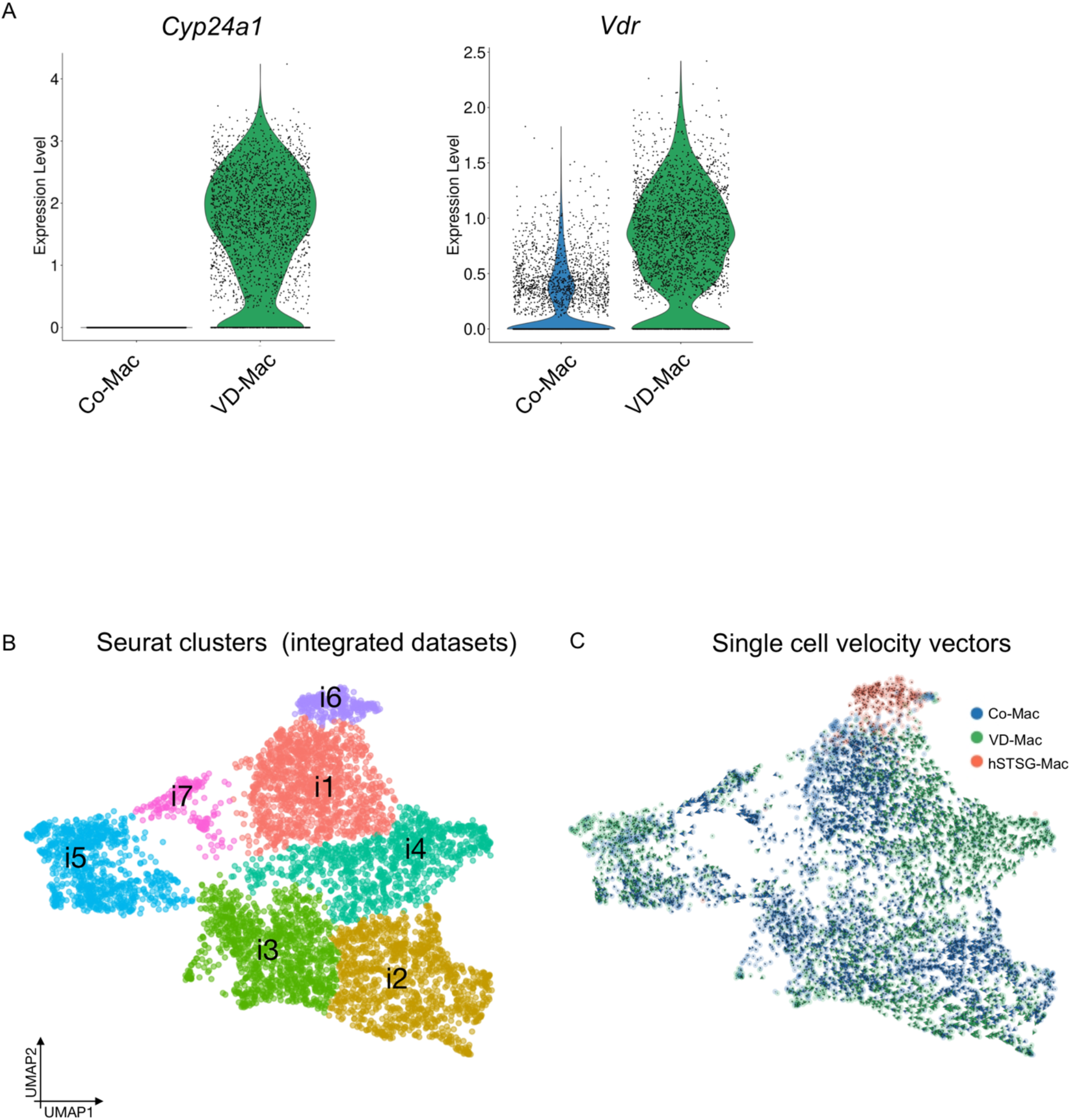
scRNAseq analysis of bone marrow derived macrophages. (**A**) Expression of *Cyp24a1* (encodes for VD3 24-hydroxylase) and *Vdr* (vitamin D receptor) in control (Co-Mac) and vitamin D3-treated macrophages (VD-Mac). (**B**) 11 distinct cell clusters were identified in the integrated dataset (cultured macrophages integrated with Trem2+ cells (cluster 0) form *in vivo* dataset). (**C**) Single cell RNA velocity vectors generated with scVelo.

**Fig. S8.**
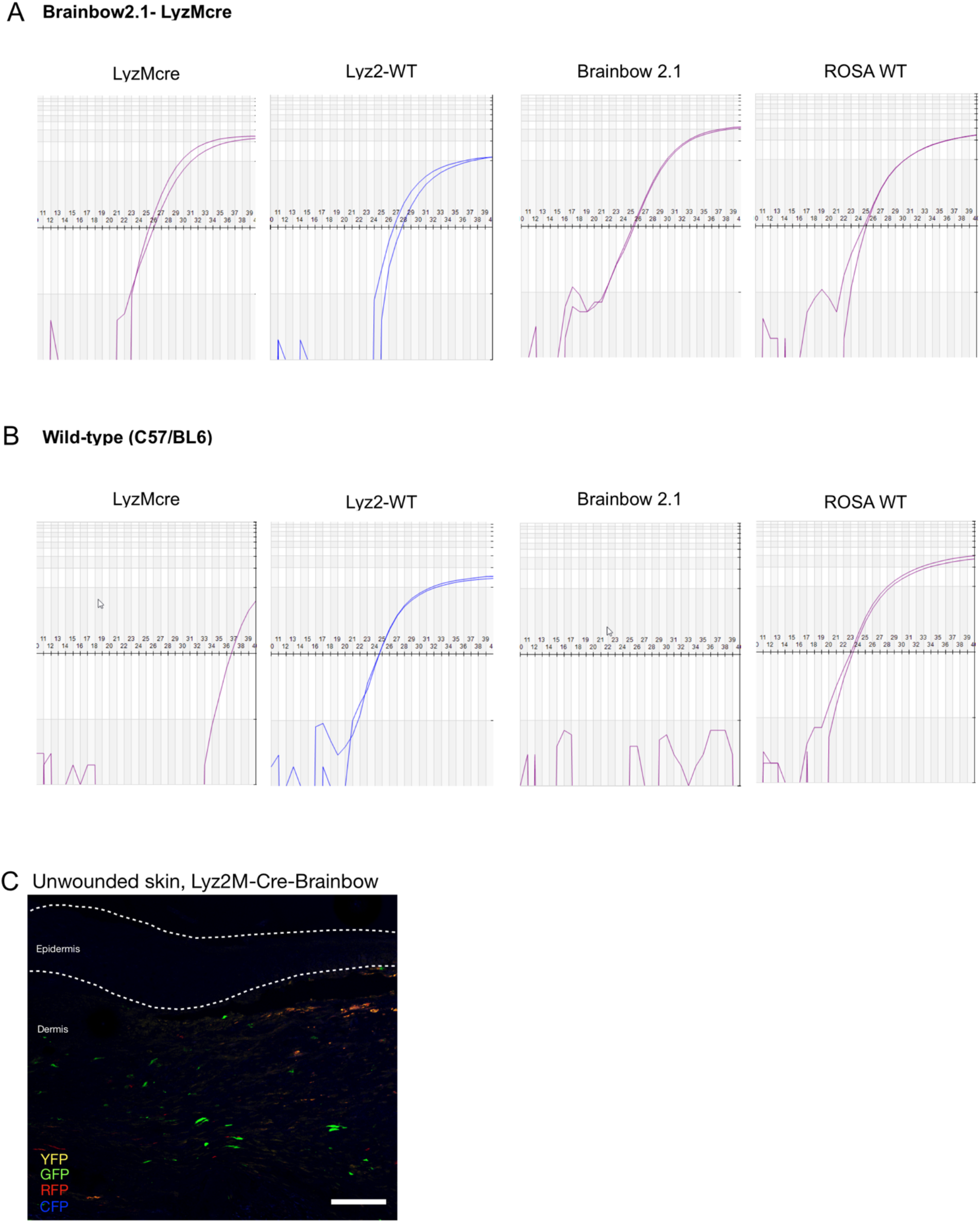
Genotype confirmation of Brainbow 2.1-LyzMcre mice and histology of unwounded skin. LyzMcre mice were crossed with R26R-Brainbow 2.1 mice. Exemplary real-time PCR (qPCR) amplification curves confirming the genotype of (**A**) Brainbow 2.1-LyzMcre mouse hemizygous for the LyzMcre and Brainbow 2.1 alleles and (**B**) C57BL6, wild-type (WT) mouse homozygous for the Lyz2 and Rosa WT alleles. X-axis = cycle threshold (CT). (**C**) Unwounded skin of Brainbow 2.1-LyzMcre mouse. Dotted line indicates the location of the epidermis, scale bar = 100 µm.

## Acknowledgements

We thank Yujin Park for her assistance in histopathology and Theresa Carlomagno for administrative support.

## Author contributions

Conceptualization: DH, KC, MTL, GCG

Methodology and investigation: DH, KC, ZNM, CAB, JAB, CJM, AHG, SEMI, DS, JQL, JP, DF, AM, DN, DCW

Data analysis: DH, KC, TF, DF, JQL, AK

Project administration: UK, DCW, MTL, GCG

Supervision: UK, MTL, GCG

Writing – original draft: DH, KC

Writing – review & editing: DH, KC, TF, JP, UK, DCW, MJ, AK, MTL, GCG

